# Mutation saturation for fitness effects at human CpG sites

**DOI:** 10.1101/2021.06.02.446661

**Authors:** Ipsita Agarwal, Molly Przeworski

## Abstract

Whole exome sequences have now been collected for millions of humans, with the related goals of identifying pathogenic mutations in patients and establishing reference repositories of data from unaffected individuals. As a result, we are approaching an important limit, in which datasets are large enough that, in the absence of natural selection, every highly mutable site will have experienced at least one mutation in the genealogical history of the sample. Here, we focus on putatively-neutral, synonymous CpG sites that are methylated in the germline and experience mutations to T at an elevated rate of ~10^-7^ per site per generation; in a sample of 390,000 individuals, ~99% of such CpG sites harbor a C/T polymorphism. These CpG sites provide a natural mutation saturation experiment for fitness effects: as we show, at current sample sizes, not seeing a polymorphism is indicative of strong selection against that mutation. We rely on this idea in order to directly identify a subset of highly deleterious CpG transitions, including ~27% of possible loss-of-function mutations, and up to 21% of possible missense mutations, depending on the type of site in which they occur. Unlike methylated CpGs, most mutation types, with rates on the order of 10^-8^ or 10^-9^, remain very far from saturation. We discuss what this contrast implies about interpreting the potential clinical relevance of mutations from their presence or absence in reference databases and for inferences about the fitness effects of new mutations.

---

A central goal of human genetics is to identify pathogenic mutations and predict how likely they are to cause disease. One approach to pinpoint sites at which mutations are likely to be deleterious is to examine whether they appear to be under purifying selection. For instance, comparisons of sequences across species have been widely used to identify highly conserved genomic regions maintained by selection over millions of years, presumably because of their functional importance (e.g., refs 1–3). The same general approach is also useful when applied within humans, where information about purifying selection is contained in whether or not a site is segregating a mutation and at what frequency (4–10).

For this application, however, the miniscule mutation rate at a typical site in the genome (~10^-8^ per base pair per generation) poses a major difficulty, as a site may be monomorphic simply by chance, i.e., when mutations at that site have no fitness consequences at all or, at the other extreme, because the mutations are embryonically lethal. Consequently, samples of hundreds or even thousands of humans, in which most sites are monomorphic, contain almost no information to distinguish strongly selected sites. In part to overcome this limitation, public repositories of human exome sequences have now grown to include data from hundreds of thousands of individuals (8,9,11–13). With these sample sizes, we should finally be able to reliably learn about fitness effects of mutations, at least at the subset of sites with elevated mutation rates.

## Mutation saturation at CpGs

To evaluate this notion, we focus on CpG sites methylated in the germline, since these are known to experience mutations much more frequently than any other type of site in the human genome (14–16); Supplementary Fig. 1). An attractive feature of methylated CpG sites is that a single mechanism, the spontaneous deamination of methyl-cytosine, is believed to underlie the uniquely high rate of C>T mutations at these sites (14); thus, germline methylation at CpG sites is strongly predictive of their mutability (16–18); Supplementary Fig. 2). We define “methylated” CpG sites in exons as those that are methylated ≥70% of the time in both testes and ovaries. For these ~1.1 million sites (of 1.8 million total CpG sites in sequenced exons), we calculate a mean haploid, autosomal C>T mutation rate of 1.17 × 10^-7^ per generation using de novo mutations in a sample of ~2900 sequenced parent-offspring trios (Methods, Supplementary Figs. 1-2; ref 19).

Although methylation levels are the dominant predictor of mutation rates at CpG sites, they are not the only one. Notably, CpG transitions differ somewhat in their mutation rates based on their trinucleotide (Supplementary Fig. 3a; ref 20); even so, they are consistently an order of magnitude higher than the genome average (16–18). Broader scale features, such as replication timing, have also been reported to shape mutation rates (21,22); nonetheless, considering methylated CpGs inside and outside exons, which differ in a number of these features, there is no appreciable difference in average DNM rates (FET p-value = 0.08, Supplementary Fig. 4a). Similarly, the rate at which two DNMs occur at the same site, a summary statistic that reflects the variance in mutation rates, is not significantly different for methylated CpGs inside versus outside exons (FET p-value = 0.5; Supplementary Fig. 4b). Thus, while there is some variation in mutability per site among methylated CpGs, it appears to be small relative to the mean mutation rate across all methylated CpGs.

Considering all such CpG sites therefore, we ask what fraction are segregating at existing sample sizes. To this end, we collate polymorphism data made public by gnomAD (9), the UK Biobank (12), and the DiscovEHR collaboration between the Regeneron Genetics Center and Geisinger Health System (11) in order to ascertain whether both C and T alleles are present in a sample of ~390K individuals (Methods).

To focus on the subset of genic changes most likely to be neutrally-evolving, we consider the ~350,000 methylated CpG sites at which C>T mutations do not change the amino acid. At these sites, 94.7% of all possible synonymous CpG transitions are observed in the gnomAD data alone, and 98.8% in the combined sample including all three datasets (Fig. 1). In other words, nearly every methylated CpG site where a mutation to T is putatively neutral has experienced at least one such mutation in the history of the sample of 390K individuals. Even in the least mutable CpG trinucleotide context, 98% of putatively neutral sites are segregating in current samples (Supplementary Fig. 3b). These observations imply that in the absence of selection, almost every methylated CpG site would be segregating a T at current sample sizes--and further that not seeing a T provides strong evidence it was removed by selection.

**Fig. 1.**
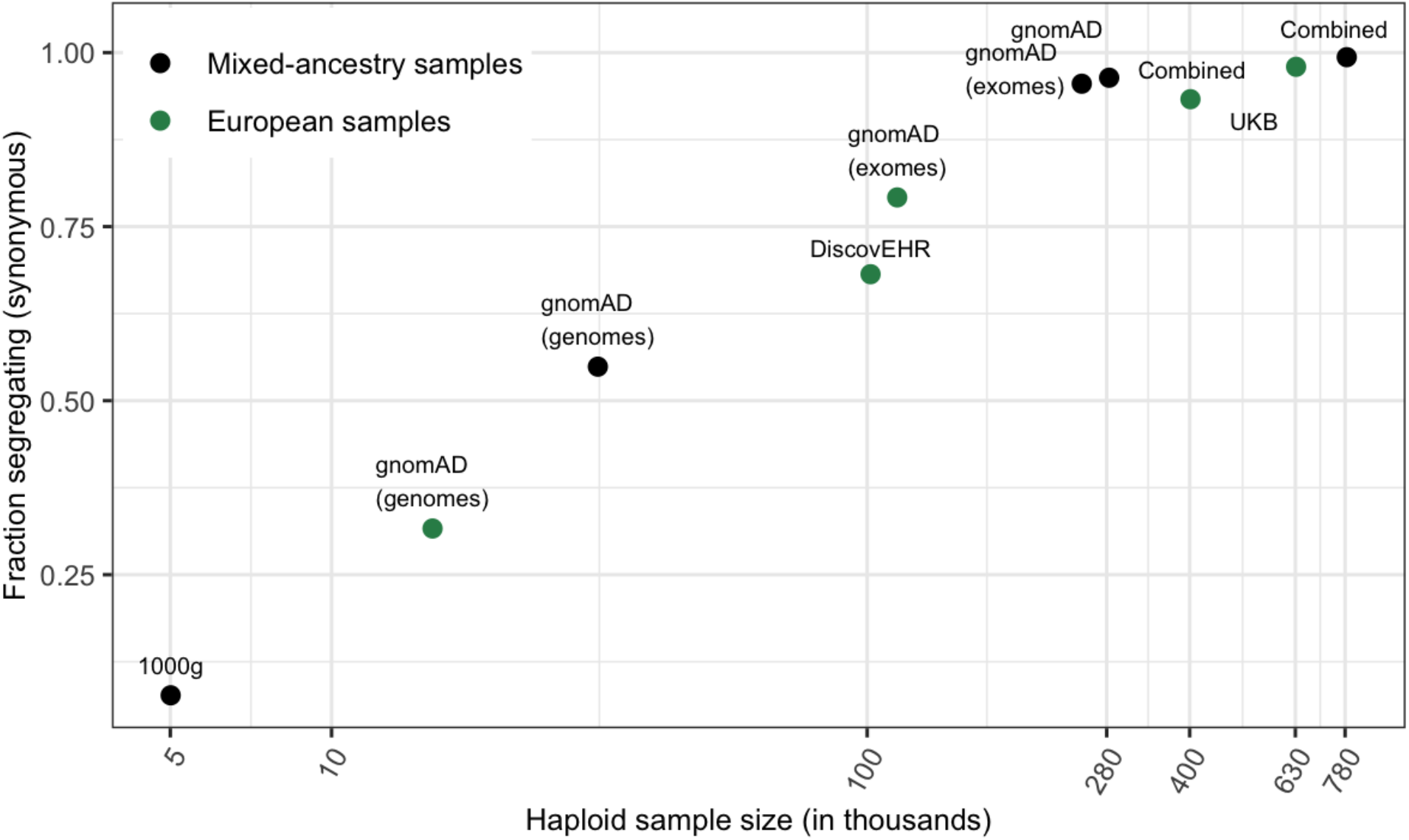
Fraction of methylated CpG sites that are polymorphic for a transition, for each sample. The combined dataset encompasses three non-overlapping data sources: gnomAD (v2.1), the UK Biobank (UKB), and the DiscovEHR cohort. “European” samples include the “Non-Finnish European” subsets of exome and whole genome datasets in gnomAD, as well as the UK Biobank and DiscovEHR, which have >90% samples labeled as of European ancestry.

## Testing a neutral model for individual sites

The mutation saturation at methylated CpG sites thus provides a robust approach to identify individual sites that are not neutrally-evolving (Supplementary Data). One way to view it is in terms of a p-value: under a null model with no selection, from which we assume that synonymous sites are drawn, all but 1.2% of neutral sites are segregating in a sample of 390K individuals. Therefore, if a given non-synonymous site, say, is invariant in a sample of ≥390K individuals, we can reject the neutral null model for this site at a significance level of 0.012.

This approach implicitly assumes that synonymous and non-synonymous sites do not differ in their distributions of mutation rates and that their distributions of genealogical histories are also the same, i.e., that the two types of sites are subject to comparable effects of linked selection. While we cannot examine whether the distributions of mutation rates are the same for lack of data, we verify that the mean de novo mutation rates do not differ for synonymous sites and for various non-synonymous annotations (Fig. 2a); we also check that the distributions of methylation levels (conditional on ≥70%), an important determinant of mutation rates, are highly similar for synonymous and non-synonymous sites (with a significant but small shift towards higher methylation and thus presumably higher mutation rates for non-synonymous sites; Supplementary Fig. 5). In turn, the standard assumption of similar distributions of genealogical histories seems sensible, given that the sites are interdigitated within genic regions (23). Under these few and at least somewhat testable assumptions, the approach based on mutation saturation at methylated CpG sites then enables us to directly pinpoint sites that are not neutrally-evolving. We note further that if synonymous sites are not all neutral and instead some fraction are under weak selection, the same idea would apply, but the null model would have to be modified accordingly.

**Fig. 2.**
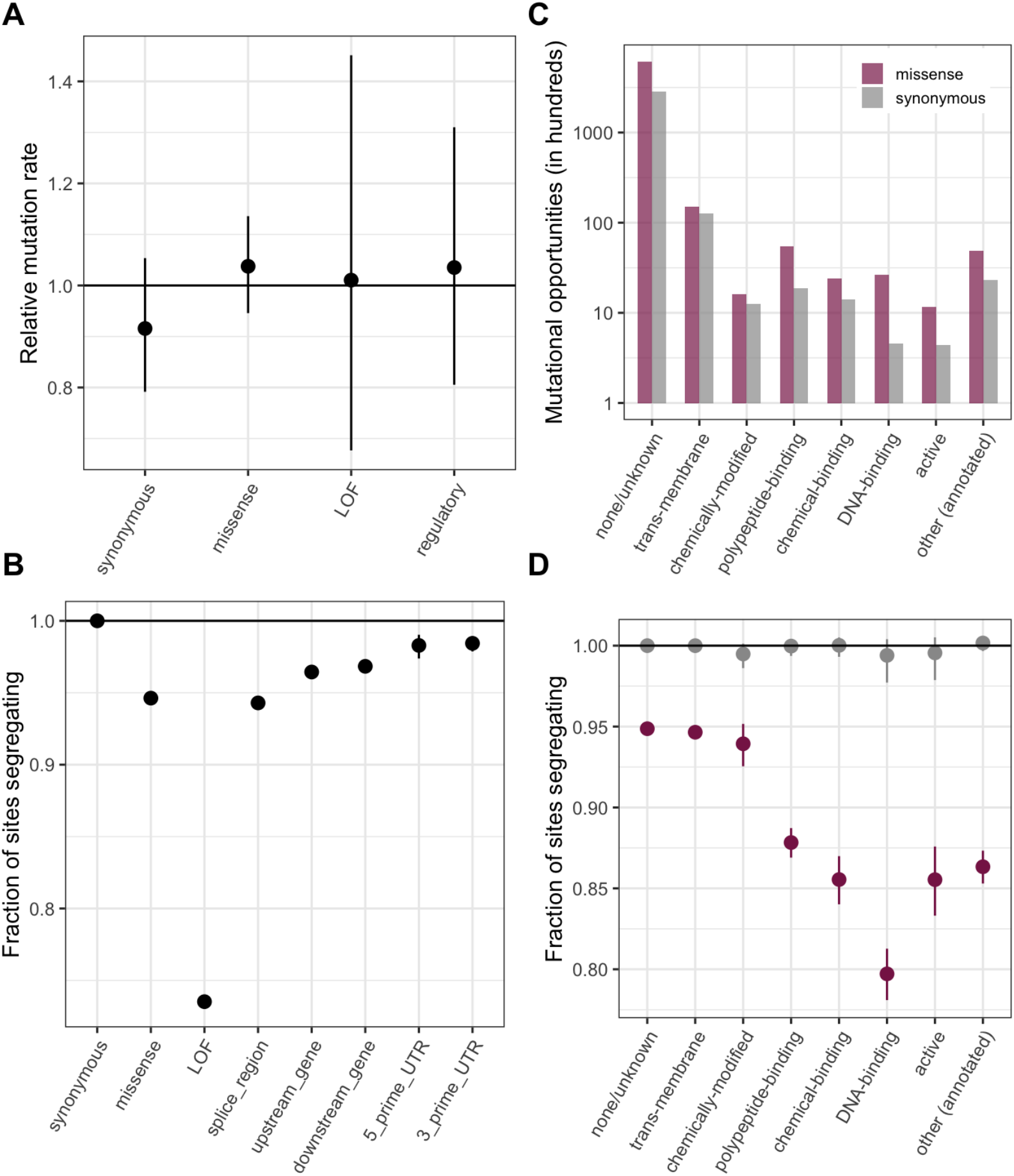
(a) DNM rates for CpG transitions at highly methylated sites by annotation class, rescaled by the total DNM rate in exons. Fisher exact tests (FETs) of the proportion of sites with DNMs in each annotation compared to all other annotations yield p-values > 0.1 in all cases. (b) Fraction of highly methylated CpG sites that are segregating as a C/T polymorphism in an annotation class, relative to the fraction of synonymous sites segregating. Error bars are 95% confidence intervals assuming the number of segregating sites is binomially distributed (FET p-values<<10^-5^ for comparisons of all annotations with synonymous sites; Methods). LOF variants are defined as stop-gained and splice donor/acceptor variants that do not fall near the end of the transcript, and meet the other criteria to be classified as “high-confidence” loss-of-function in the gnomAD data. (c) The amount of data for synonymous and missense changes involving highly methylated CpG transitions by the type of functional protein site. (d) The proportion of synonymous and missense segregating C/T polymorphisms in different classes of functional sites. Error bars are 95% confidence intervals assuming the number of segregating sites is binomially distributed (FET p-values<<10^-5^ for comparisons of all missense annotations with synonymous sites; Methods). All annotations are obtained using the canonical transcripts of protein coding genes (see Methods).

## Comparing the fraction of segregating sites across annotations

Under these same weak assumptions, it is also possible to compare the proportion of methylated CpG sites segregating a transition across annotations. Here, we consider the fraction of sites segregating a transition in each annotation class in a sample of 780k chromosomes, rescaled by the fraction segregating at synonymous sites. All categories of missense, loss-of-function, and regulatory variants show a significant depletion in the fraction of segregating sites compared to synonymous variants (Fig. 2b). The deficit for a given annotation, due to selection, is an indicator of the average deleteriousness of de novo mutations in that annotation.

Notably, these data suggest that there are ~27% fewer loss-of-function variants than would be expected under neutrality. Supporting the widely-used assumption that true LOF mutations within a gene are equivalent (after filtering for those at the end of transcripts; e.g., refs 9, 24), when we compare the set of CpG sites at which mutations are annotated as leading to protein-truncation in the first versus the second half of transcripts, approximately the same number are missing relative to synonymous sites in both subsets (Supplementary Fig. 6; FET p-value =0.09). A 27% deficit of loss-of-function variants is again seen if we match the sites to synonymous mutational opportunities with the same predicted level of linked selection, i.e., with similar genealogical histories (ref 25; Supplementary Fig. 7a).

By comparison, the fraction of missense mutations and splice region mutations not observed in current samples is only about 6% (whether or not we match for the effects of linked selection; see Supplementary Fig. 7a). While LOF and missense annotation classes are most commonly used in determinations of variant pathogenicity, any two sets of methylated CpGs with similarly-distributed mutation rates can be ranked in this manner. As one example, we stratify missense mutations by the type of functional site in which they occur. Strikingly, for the subset of sites at which missense mutations may disrupt or alter binding, particularly DNA-binding, there is a ~21% deficit in segregating sites relative to what is seen at synonymous sites, in contrast, say, to the much smaller deficit at missense changes within trans-membrane regions (Fig 2c-d, Supplementary Fig. 8; Supplementary Fig. 7b).

A widely-used alternative to the use of annotations is to stratify sites by their predicted functional importance with scores such as CADD (26). Mean transition rates at methylated CpGs are similar across CADD deciles, as expected if there is limited variation across them (and DNMs are rarely embryonic lethal) (Supplementary Fig. 9a). 17% of methylated CpG sites are monomorphic for the top decile of CADD scores and as expected, the fraction of segregating sites decreases with increasing CADD scores, confirming that, when considering sets of sites with the same mutation rate, the observed deficit in segregating sites provides a ranking of the sets by the typical deleteriousness of mutations (Supplementary Fig. 9b).

Considering a more heterogeneous set of sites than methylated CpGs, however, such as all CpG sites in exons, we can no longer assume mutation rates to be the same across CADD deciles, and indeed when we examine the mean transition DNM rate, it varies significantly. One implication is that these scores, while meant to isolate the effects of selection, may nonetheless confound mutation rates and conservation to some extent. In particular, among relatively high CADD scores in the genome are likely some sites that are not in fact unusually constrained, but have a lower mutation rate.

## What can be learned about other mutation types?

Given that current exome samples are informative about selection on transitions at methylated CpGs, a natural question is to ask to what extent there is also information for less mutable types, with mutation rates on the order of 10^-8^ or 10^-9^ per site per generation. For sites with mutation rate on the order of 10^-9^, which is the case for the vast majority of non-CpGs, the fraction of possible synonymous sites that segregate in a sample of 780K chromosomes is very low: for instance, it is ~4% for T>A mutations, which occur at an average rate of 1.2×10^-9^ (Supplementary Fig. 1) and ~30% even for other C>T mutations, which occur at a rate of 0.9×10^-8^ per site (Supplementary Fig. 1), compared to ~99% for C>T mutations at methylated CpGs (Fig. 3a). For invariant sites of these less mutable types, there is little information with which to evaluate the fit to the neutral null in current samples. Reflecting this lack of information, in the p-value formulation, monomorphic sites would be assigned p≤0.96 for T>A sites and p≤0.7 for C>T sites.

**Fig. 3.**
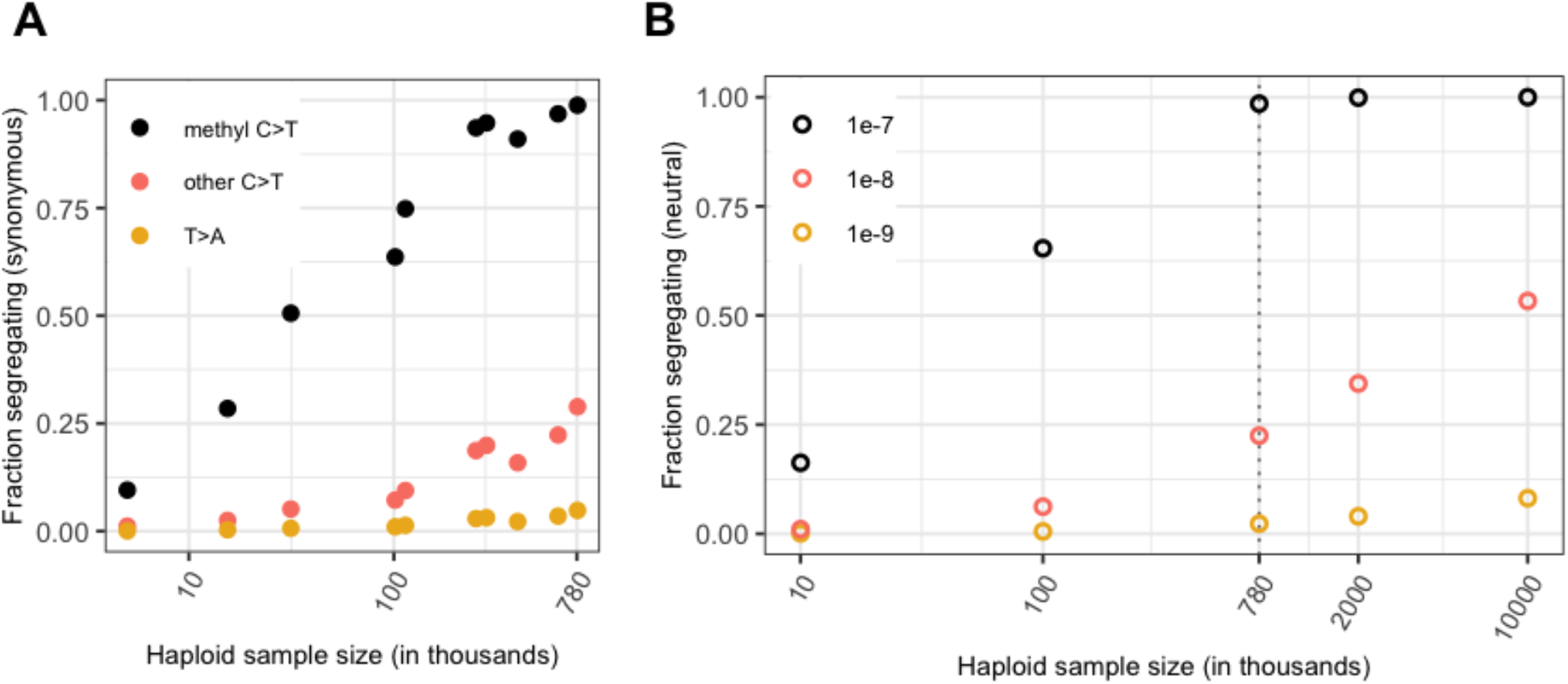
(a) Fraction of possible synonymous C>T mutations at CpG sites methylated in the germline and at all other C sites, and the fraction of possible synonymous T>A mutations that are observed in a sample of given size. (b) Probability of a polymorphism in simulations, assuming neutrality, a specific demographic model and a given mutation rate (see Methods).

How large samples have to be for other mutation types to reach saturation depends on the length of the genealogy that relates sampled individuals, i.e., the sum of the branch lengths, which corresponds to the number of generations over which mutations could have arisen at the site. For a mutation that occurs at rate 10^-9^ per generation, the average length of the genealogy such that exactly one mutation would be expected is 10^9^ generations. That synonymous CpG sites are close to saturation when they experience mutations to T at a rate of of 1.17 × 10^-7^ per generation implies that the average length of the genealogy relating the 390K individuals is at least 8.5 million (1 divided by 1.17 × 10^-7^) generations--likely quite a bit longer given that more than one mutation has occurred at a substantial fraction of sites (9,27). On the other hand, the observation that mutation types with rates on the order of 10^-8^ are far from saturation indicates that the average length of the genealogy for these 390K individuals is substantially shorter than 100 million generations. The length of the genealogy is expected to increase much more slowly than linearly with the number of samples (28); thus, saturation under a mutation rate of 10^-9^ per site per generation may not even be achievable in samples ten times larger than at present.

To explicitly examine the relationship between sample size, mutation rate and the amount of variation at a locus, we simulate neutral evolution at a single site with the three different mutation rates above, under a variant of the widely-used Schiffels-Durbin demographic model for population growth in Europe (29), in which we set the effective population size *N_e_* equal to 10 million for the past 50 generations (Methods). While this model is clearly an oversimplification, it recapitulates what we see in data for synonymous mutations for different mutation types reasonably well (Fig. 3). Consistent with the rough estimate above, under our choice of demographic model, a sample of 780K chromosomes has a genealogy spanning an average of 34 million generations (Supplementary Fig. 10a). Increasing the number of samples by a factor of 12 only increases the average tree length ~3.3x (Fig. 3b, Supplementary Fig. 10a); thus, a site that mutates at rate 10^-9^ per generation is expected to have experienced ~0.04 mutations in the genealogical history of a sample of ~1 million, and 0.1 mutations in a sample of 10 million.

Quantitative predictions of our model are not reliable, given uncertainty about the demographic history and in particular the recent effective population size in humans (Supplementary Fig. 10b). Moreover, for simplicity, we model one or at most two populations, when mixed ancestry samples have slightly longer genealogical histories (Supplementary Fig. 10c); thus, as samples diversify by ancestry, more variation will be captured than if only individuals from similar ancestries are included. Perhaps most importantly, for the very large sample sizes considered here, the multiple merger coalescent is a more appropriate model (30). Nonetheless, the qualitative statement that less mutable types will remain very far from saturation in the foreseeable future should hold.

In the absence of information about single sites for most mutational types in the genome, it is still possible to learn to a limited degree about selection using bins of sites. If we construct a bin of *K* synonymous sites with the same average mutation rate per bin as a single methylated CpG, then at least one site per bin is polymorphic in ~99% of bins (see Supplementary Fig. 11 for an example with T>A mutations and *K*~100), just as ~99% of individual methylated CpG sites are segregating. Thus, if a bin of *K* non-synonymous sites with the same average mutation rate is invariant, the p-value associated with the bin is 0.01, indicating that one or more sites in the bin is unlikely to be neutrally-evolving.

## How strong is the selection that leads to invariant methylated CpG sites?

Leveraging saturation to identify a subset of sites that are not neutrally-evolving makes appealingly few assumptions, but provides no information about how strong selection is at those sites. To learn about the strength of selection consistent with methylated CpG sites being monomorphic, a series of strong assumptions are needed: we require a demographic model, a prior distribution on *hs* and a mutation rate distribution across sites. Here, we assume a relatively uninformative log-uniform prior on the selection coefficient *s* ranging from 10^-7^ to 1 and fix the dominance coefficient *h*=0.5 (as for autosomal mutations with fitness effects in heterozygotes, we only need to specify the compound parameter *hs*; reviewed in ref 31), as well as a fixed mutation rate of 1.2×10^-7^ per site per generation. We rely on the demographic model for population growth in Europe described above (29); as is standard (e.g., refs 4,6,7,24,32–34), we also assume that *hs* is fixed over time, even as the effective population size changes dramatically. Under these assumptions, we estimate the posterior distribution of *hs* at a site, given that the site is monomorphic, segregating with 10 or fewer derived copies of the T allele, or segregating with more than 10 copies (Fig. 4a and 4b, Methods).

**Fig. 4.**
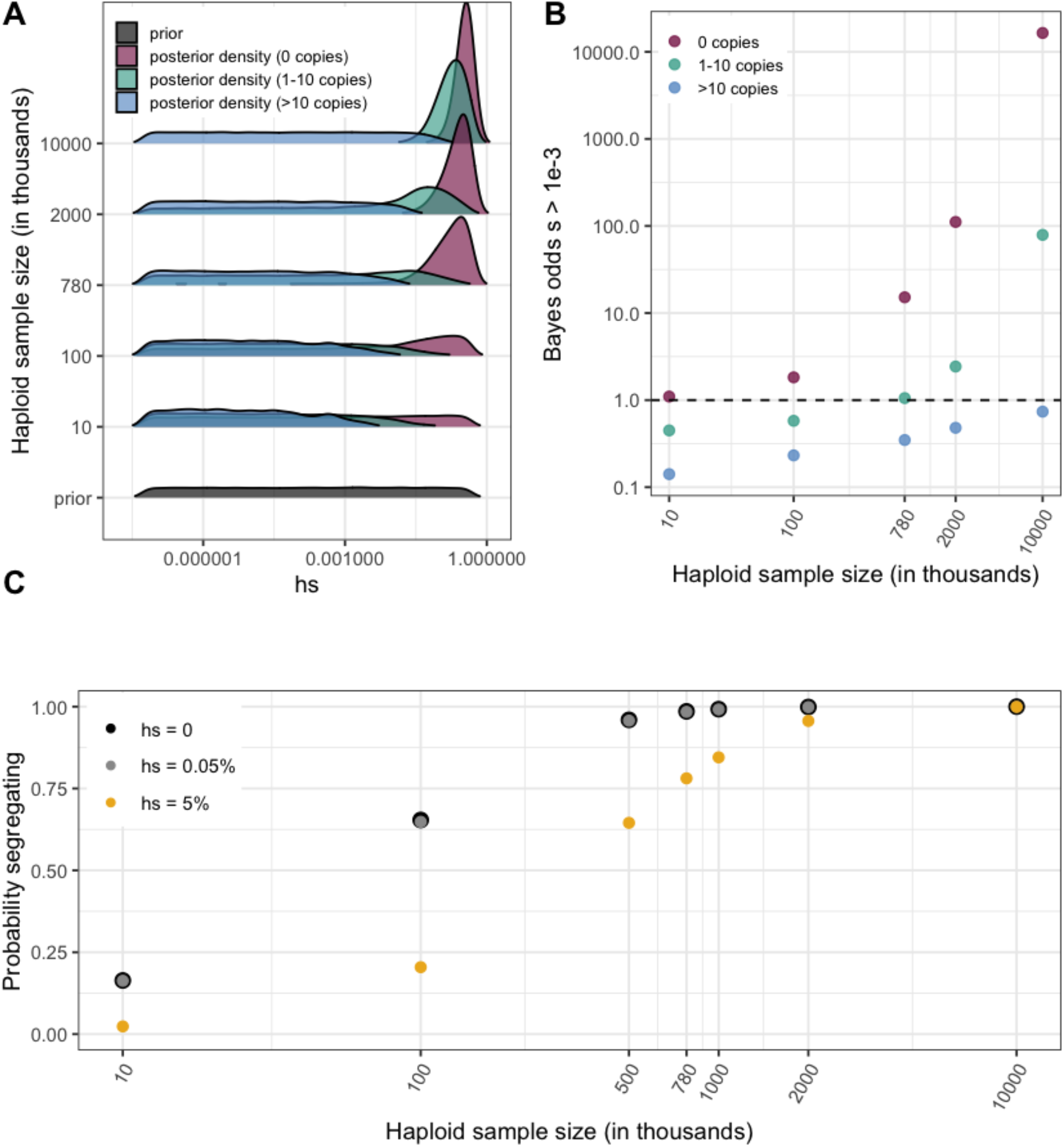
(a) Prior and Posterior log densities for *hs* for a C>T mutation at a methylated CpG site observed at 0, 1-10, or >10 copies at various sample sizes. (b) Bayes odds (i.e., posterior odds divided by prior odds) of *s* > 0.001 for a C>T mutation at a methylated CpG site observed at 0, 1-10, or >10 copies, at various sample sizes. (c) Probability of a methylated CpG site segregating a T allele in simulations, if the mutation has no fitness effects (*hs*=0) and if it is deleterious (with a heterozygote selection coefficient *hs*=0.05%) or highly deleterious (with a heterozygote selection coefficient *hs*=5%, where *s* is the selection coefficient and *h* is set to 0.5).

In small samples, in which most sites are monomorphic, being monomorphic is consistent with both neutrality and very strong selection (Fig. 4a) and the Bayes odds are close to 1 (Fig. 4b). In contrast, with larger sample sizes, in which putatively neutral CpG sites reach saturation, the posterior distribution for invariant sites is highly peaked: what is not segregating is likely strongly deleterious. At current sample sizes of 390K individuals, under our model, an invariant methylated CpG is very likely to have an *hs* > 0.5 × 10^-3^ (Fig. 4b), a conclusion that is relatively insensitive to the choice of prior (Supplementary Fig. 12).

While the relationship of selection strengths to clinical pathogenicity is not straight-forward, selection coefficients on that order are likely to be of relevance to determinations of pathogenicity in clinical settings (refs 24, 35; Agarwal, Fuller, Przeworski, in prep.). Indeed, mutations with *hs* > 0.5 x10^-3^ may be highly deleterious to some individuals that carry them, enough to produce clinically visible effects, but vary substantially in their penetrance. Our analysis suggests that the ~27% of LOF mutations and ~6% of missense mutations not seen in current samples are subject to this degree of selection.

## Interpreting monomorphic and polymorphic sites in current reference databases

Although sites that are invariant in large samples are enriched for the strongest deleterious effects, in a sufficiently large sample, even a segregating site can be subject to strong selection (Fig. 4a, 4b). For instance, in current exome sample sizes, a C>T mutation at a methylated CpG site with *hs*=0.5 × 10^-3^ is almost always observed segregating (Fig. 4c). This follows from the expectation under mutation-selection-drift balance (36): for example, in a constant population size, a mutation that arises at rate 1.2 × 10^-7^ per generation and is removed by selection at rate *hs*=0.05% per generation is expected in the population at frequency 2.4 × 10^-4^ on average; in a sample of 780k, the mean number of copies is 187. Even with substantial variation due to genetic drift and sampling error, such a site should almost always be segregating at that sample size. In fact, even a mutation with *hs* of 5% would quite often be observed. Thus, segregating sites in large samples are a mixture of neutral, weakly selected and strongly selected sites. An implication is that, although large reference repositories such as gnomAD were partly motivated by the possibility of excluding deleterious variants--with the idea that seeing a variant of unknown function in a reference data set is suggestive that the variant is benign--as samples grow in size, it cannot simply be assumed that clinically relevant variants are absent from reference datasets. In principle, the only mutations never seen as samples grow in size would be the ones that are embryonically lethal.

More generally, any interpretation of variants of unknown function by reference to repositories such as gnomAD or disease cohorts enriched for deleterious variation (e.g., ref 37), whether the goal is to exclude benign variants or identify likely pathogenic ones, is implicitly reliant on assumptions that change with sample size and dramatically differ by mutation type. At current sample sizes, invariant methylated CpGs are likely highly deleterious; for less mutable types, the information content at invariant sites is negligible at even the largest sample sizes considered (Supplementary Fig. 13). Similarly, learning about the fitness consequences of mutations from their observed frequencies is contingent on assumptions about the mutation rate, and the demographic history of the sample.

## The distribution of fitness effects in the human genome

Modeling the distribution of fitness effects (DFE) has a long history in population genetics (4,5,38), but until recently, inferences were based on genetic variation in samples of at most a couple of thousand chromosomes (e.g., refs 5–7,32,33). As is well appreciated, the fitness effects at the few sites segregating in such samples are a small and biased draw from the DFE and thus the inferred distribution of fitness effects is unlikely to recapitulate the true DFE in the genome. Moreover, for lack of sufficient information with which to distinguish weakly from strongly selected mutations, a number of approaches have relied on a specific and arbitrary parametric form for the distribution of fitness effects across sites. In that regard, not only do inferences based on small samples result in relatively noisy parameter estimates, the results can be misleading, especially about the fraction of sites under strong selection (5–7,32,33).

Because current samples in humans are vastly more informative, these limitations can now start to be circumvented. As we show, a typical site in the genome, with a mutation rate of 10^-8^ per generation, does not provide much information about selection (Supp Fig 13), because the average length of the genealogy is perhaps on the order of 10^7^ generations. One exception, which is a special case, is gene loss: each gene can be conceived of as a single locus at which many possible LOF mutations have the same fitness impact (refs 9,24,31,39; Agarwal, Fuller, Przeworski, in prep.). The mutation rate to LOF, calculated by summing rates of individual LOF mutations, is ~10^-6^ per gene per generation on average (9), such that in the absence of selection, many LOF mutations are expected in most genes. Another way to overcome current sample size limitations might be to bin sites and perform inferences on the bin; however, if mutations at sites within a bin vary in their fitness effects, inferences based on these bins are less straight-forward, as the mutation frequency in a bin reflects the harmonic mean of *hs* across sites in the bin (see Methods).

Given these limitations, individual methylated CpG sites can provide a useful point of entry to understanding the DFE in humans. Although methylated CpG sites appear under somewhat less constraint than other sites, the differences are subtle (Supplementary Fig. 14), and what we learn at these sites can tell us what to expect more generally. In current samples, the posterior odds for invariant methylated CpGs having *hs* ≥ 5×10^-4^ are 92% under our model, whereas they are 36% for segregating methylated CpGs. Considering only possible LOF mutations at methylated CpGs, of which 27% are not observed in current samples, this implies that the fraction of LOF mutations with this degree of selection is roughly 51% (=0.27×0.92+0.73×0.36). Although crude, this estimate is remarkably close to the 53% of possible LOF variants that are found in genes that have *hs* ≥ 5×10^-4^, when *hs* is estimated for the loss-of-function of each gene using allele frequencies of all LOF mutations in gnomAD (Agarwal, Fuller, Przeworski, in prep.). This close agreement lends further support to the notion that fitness effects at methylated CpGs are informative about those at other sites, at least for some annotations.

For missense sites, given the same uninformative prior on *hs* as for LOF mutational opportunities, the fraction estimated to be highly deleterious is close to 40% (=0.06×0.92+0.94×0.36). Since on average, 0.97% of all de novo point mutations are missense and 0.06% lead to a LOF (see Methods), current estimates translate into odds of approximately 1 in 240 (=40%x0.97%+51%x0.06%) that such a mutation has an effect *hs* ≥5×10^-4^. Assuming each individual inherits roughly 70 new mutations (16,17), these estimates imply that at least one out of every ~3.4 individuals is born with a new and potentially highly deleterious, non-synonymous mutation (more if other mutation types are considered).

Moving forward, we should soon have substantial information not only about the DFE but the strength of selection at individual, methylated CpG sites (Fig. 4). Inferences based on them, or indeed any sites, will need to rely on an accurate demographic model, particularly for the recent past of most relevance for deleterious mutations; this problem should be surmountable, given the tremendous recent progress in our reconstruction of population structure and changes in humans (e.g., refs 29,40,41). Inferences will also require a good characterization of mutation rate variation across methylated CpG sites, as is emerging from human pedigree studies and other sources (e.g., refs 17,42,43). Putting these elements together, robust inference of the fitness effects of mutations in human genes is within reach, at least through the lens of CpG sites.

## Acknowledgements

We thank Peter Andolfatto, Kelley Harris, Hakhamanesh Mostafavi, Magnus Nordborg, Itsik Pe’er, Jonathan Pritchard, Guy Sella, as well as Arbel Harpak, Zach Fuller, and other members of the Andolfatto, Przeworski and Sella labs for helpful discussions. This work was supported by NIH grants GM121372 and GM122975 to MP.

## Methods

### 1. Processing de novo mutation data

We focused on ~190,000 published de novo mutations in a sample of 2976 parent-offspring trios that were whole genome sequenced (19). To date, this is the largest publicly available set of trios that, to our knowledge, have not been sampled on the basis of a disease phenotype. Unless otherwise specified, we used these DNMs to calculate mutation rates, as described in later sections. We converted hg38 coordinates to hg19 coordinates using UCSC Liftover. We excluded indels, and all DNMs that occur outside the ~2.8 billion sites covered by gnomAD v2.1.1 whole genome sequences. We obtained the immediately adjacent 5’ and 3’ bases at each position from the hg19 reference genome, so that we had each de novo mutation within its trinucleotide context; we used this information to identify CpG sites. Where such data were available (for 89% of CpG de novo mutations), we also annotated each site with its methylation status measured by bisulfite sequencing in testis sperm cells and ovaries (see Supplementary Table 1).

We annotated DNMs with their variant consequences using Variant effect predictor (v87, Gencode V19) and the hg19 LOFTEE tool (9) to flag high-confidence (“HC”) loss-of-function variants. We obtained the fraction of DNMs in the genome that occured at sites annotated as missense or LOF in the “canonical” protein-coding transcript for each gene provided by Gencode.

### 2. Processing polymorphism data

We downloaded publicly available polymorphism data from gnomAD (9), the UK Biobank (12), the DiscovEHR collaboration between the Regeneron Genetics Center and Geisinger Health System (11), and 1000 Genomes Phase 3 (44). Where needed, we lifted over coordinates to the hg19 reference assembly using the UCSC LiftOver tool. Salient characteristics of these samples are summarized below.

| Dataset | Regions Sequenced | Individuals | Variants | Populations sampled | Original alignment |
| --- | --- | --- | --- | --- | --- |
| 1000 genomes Phase 3 (also included in gnomAD) | Genomes | 2,504 | 84 million | mixture | hg19-b37 |
| gnomAD v2.1.1 | Exomes | 125,748 | 15 million | mixture | hg19 |
| gnomAD v2.1.1 | Genomes | 15,708 | 230 million | mixture | hg19 |
| UK Biobank | Exomes | 200,643 | 16 million | ~93% European ancestry | hg38 |
| DiscovEHR | Exomes | 50,726 | 8 million | ~98% European ancestry | hg19-b37 |

For the gnomAD data, we obtained the allele frequency for each variant in the full exome and genome samples, as well as their Non-Finnish European (“NFE”) subsets from the VCF files (in hg19 coordinates) provided. For each sample, we obtained the set of segregating sites (i.e., the set of variants that pass gnomAD quality filters and have an allele frequency > 0 in the sample). For the 1000 Genomes Phase-3 data, we obtained the set of variant positions similarly. Note that the 1000 Genomes samples are also contained within the gnomAD sample. For the DiscovEHR sample, allele frequencies are available where MAF > 0.001 (and set equal to 0.001 for lower values>0); this information allows us to determine the set of sites segregating in this sample, but we do not have access to any other information about individual variants.

For the UK Biobank exome sequencing data, additional processing was required. We downloaded the population-level plink files with exome-wide genotype information for ~200,000 individuals. We excluded exome samples that did not pass variant or sample quality control criteria in the previously released genotyping array data. Specifically, we excluded samples that have a discrepancy between reported sex and inferred sex from genotype data, a large number of close relatives in the database, or are outliers based on heterozygosity and missing rate, as detailed in Bycroft et al., 2018 (45). Finally, we excluded individuals who withdrew from the UK Biobank by the end of 2020. This left us with 199,930 exome samples that overlap with the high-quality subset of the genotyped samples. We additionally limited our analysis to the list of ~40 million exonic sites with an average of 20x sequence coverage provided by UK Biobank, and for which variants met the QC criteria described in Szustakowski et al., 2020. We transformed the processed plink files into the standard variant call format, polarized variants to the hg38 reference assembly, and obtained the frequency of the non-reference allele in the sample. We then lifted over the coordinates from hg38 to hg19 using the UCSC LiftOver tool. We excluded the few positions where the reference alleles were mismatched or swapped between the two assemblies.

### 3. Identifying and annotating mutational opportunities in the exome

For all possible mutational opportunities in sequenced exons, we collated a variety of functional annotations. To this end, we first generated a list of all possible SNV mutational opportunities in the exome. We obtained the list of sites that fall in exons or within 50 base pairs (bp) of exons in Gencode v19 genes and that are among the ~2.8 billion sites covered by gnomAD v2.1.1 whole genome sequences. For each position, we extracted the reference allele from the hg19 assembly and generated the three possible single-nucleotide derived alleles. We also obtained the immediately adjacent 5’ and 3’ bases at each position from the hg19 reference genome, so that we had each mutational opportunity within its trinucleotide context; we used this information to identify CpG sites. Where such data were available, we also annotated each site with its methylation status in testis sperm cells and ovaries.

To identify sites at which variants or de novo mutations could be confidently assayed by whole-exome sequencing methods, we obtained regions targeted in whole exome sequencing from gnomAD and the UK Biobank. We limited our analysis to sites that were covered at 20x or more in the exome sequencing subsets of both gnomAD and UK Biobank (that lifted over correctly to the hg19 assembly), which we refer to as “accessible sites”.

We then annotated the ~90 million mutational opportunities (at 30 million sites) with CADD scores and variant consequences using Variant effect predictor (v87, Gencode V19) and the hg19 LOFTEE tool (9) to flag high-confidence (“HC”) loss-of-function variants. For loss-of-function variants, we also noted their location in the gene by exon number (e.g., in exon 10 of 12 exons in the gene). We used a database of curated protein features derived from Refseq proteins (46) to annotate all sites in protein coding genes that were associated with a particular type of functional activity (detailed functional annotations were available for 62,387 of 1.1 million methylated CpG sites). At each site, we used either the primary “site-type” annotation, or when that was missing or listed as “other”, we extracted the annotation from the more detailed “notes” field where this information was provided.

Because there are multiple transcripts for each variant, we limited our analysis to the “canonical” protein-coding transcript for each gene provided by Gencode to obtain a single annotation for each variant. For 10-20% of variants, this approach still yielded multiple possible consequences per variant, for instance, where there are multiple canonical transcripts due to overlapping genes. For these cases, we assigned one of the “canonical” transcripts to the variant at random, to avoid making assumptions about their relative importance. Further overlaps within the same gene, e.g., a missense variant that is also a splice variant in the same transcript, or a DNA-binding site that also undergoes a particular post-translational modification were resolved in the same manner.

As an alternative approach, we obtained the worst consequence in all protein-coding transcripts for each variant, using the ranks of variant consequences by severity provided by Ensembl (see Supplementary Table 1). In the absence of systematic ranking criteria for the protein function annotations we used the following order: sites that were designated as having catalytic activity (“active” sites) were given highest priority in overlaps, followed by DNA-binding sites, followed by other types of binding (to metal, polypeptides, ions), and finally by sites that are known to undergo post-translational or other regulatory modifications, and trans-membrane sites. Thus, a transmembrane site with regulatory activity is classified as a regulatory site, while a regulatory site with DNA-binding activity is classified as DNA-binding. Using these alternate criteria to group sites does not affect our conclusions (Supplementary Fig. 7).

All sources of annotation data are listed in Supplementary Table 1. A list of CpG sites and annotations is available in Supplementary Data 1.

### 4. Comparing fitness effects across sets of mutational opportunities

To assess whether the set of 1.1 million C>T mutational opportunities at methylated CpG sites are systematically different from the other ~90 million exonic mutational opportunities in their potential fitness effects, we compared the distribution of CADD scores in the two groups using a Kolmogorov-Smirnov test. We note that this comparison is likely to be somewhat confounded by differences in mutation rates, given our finding that CADD scores do not perfectly isolate the effects of selection from those of variability in mutation rates (Supplementary Fig. 8b). Since the mutation rate for methylated CpG sites is higher than for other types, this may lead them to appear somewhat less constrained than they actually are.

We further compared the fraction of C>T mutational opportunities at methylated CpGs in an annotation class vs. the fraction of other mutational opportunities in that class. We used a Fisher exact test (with a Bonferroni correction for four tests) to determine whether the two sets of mutational opportunities were differently distributed across synonymous, missense, regulatory, and LOF variant classes.

### 5. Obtaining mean de novo mutation rates by mutation type and annotation

We counted the total number of de novo mutations in sequenced exons (~91 million mutational opportunities) for 8 classes of mutations: two transitions and a transversion each at C and T sites, transitions at CpG sites with relatively low levels of methylation (defined here as < 70%), and transitions at CpG sites with high levels of methylation (≥70%). To obtain the mutation rate per site per generation, we divided the counts by the haploid sample size (2 × 2976 individuals; see section 1) and the number of mutational opportunities of each type. We report 95% confidence intervals assuming a Poisson distribution for mutation counts. The rates obtained (Supplementary Fig. 1a) are similar to previous ones (16–18) and roughly consistent with the rates predicted by the gnomAD mutation model (9).

To evaluate the impact of methylation status on the mutation rate at CpG sites, we obtained the mean mutation rate for C>T mutations at CpG sites in each methylation bin as described above, separately for methylation levels in ovaries and testes. While there is a limited amount of data, especially for some low-methylation bins, our choice of cutoff for “methylated” seems sensible (Supplementary Fig. 2).

We then calculated the mean mutation rate for methylated CpG transitions, for different compartments in the genome, namely in (a) exons and non-exons, (b) four variant consequence categories: synonymous, missense, regulatory, and LOF variants, (c) CADD score deciles, and (d) in exons that constitute the first half vs the second half of genes. We also calculated the mean mutation rate for methylated CpG transitions in four trinucleotide contexts (ACG, CCG, GCG, and TCG). In each case, we obtained the total number of de novo mutations and the Poisson 95% confidence interval around mutation counts in each group, and divided by the number of mutational opportunities in the group. We tested if the proportion of methylated CpG sites with de novo C>T mutations in each non-synonymous compartment was different from the proportion of synonymous methylated CpGs with de novo C>T mutations, accounting for multiple tests.

### 6. Variance in mutation rate at methylated CpGs

Although current samples of DNM data are large enough to compare the mean mutation rate at methylated CpGs across the annotation classes examined here, there is not enough data to directly compare variances in mutation rates. To learn how much broad scale features beyond methylation and the immediate trinucleotide context shape variation in mutation rates at methylated CpGs, we therefore rely on a broader set of regions e.g., those that fall inside and outside exons. Exonic and non-exonic regions differ considerably in epigenetic features and replication timing (21); yet, there is no discernable difference in average de novo mutation rates at methylated CpGs inside and outside sequenced exons (FET p-value = 0.08, Supplementary Fig. 4a). We also compared the number of double and single de novo hits in exons and non-exons using a Fisher exact test (p-value = 0.5, Supplementary Fig. 4b). Since the number of double hits reflects the variance in mutation rates across sites, these results lend some support to there being limited variation due to broad scale genomic features in transition rates at methylated CpGs.

### 7. Calculating the fraction of sites segregating by annotation

For each methylated CpG site in the exome, there are three mutational opportunities (C>A, C>G, C>T); we focus only on the opportunities for C>T mutations. For each methylated CpG site then, we noted whether or not it was segregating, or in other words if there was a C>T variant in samples of individuals from gnomAD (9), the UK Biobank (12), the DiscovEHR dataset (11), and 1000 Genomes Phase 3 (44), processed as described above, or a combined sample of 390K non-overlapping individuals.

Within the set of methylated CpG sites where C>T mutations are synonymous, we calculated the fraction segregating in each sample of interest. Similarly, for different subsets of methylated CpGs, namely those in (a) four variant consequence categories: synonymous, missense, regulatory, and LOF variants, (c) CADD score deciles, (d) functional site categories (e.g., trans-membrane vs catalytic sites in proteins), and (e) the first half vs the second half of genes, we calculated the fraction segregating in the combined sample of 390K individuals. We rescaled the fraction of sites segregating in each annotation by the fraction of synonymous sites segregating in the sample.

We verified that the differences in the fraction of sites segregating across annotations are not due to variable impacts of linked selection across annotations. To do so, we calculated the fraction of sites segregating with sites in different annotations matched for B-statistics (25); we obtained very similar results with this approach (Supplementary Fig. 7).

We assumed that conditional on the number of mutational opportunities and a fixed probability of segregating for each site in a compartment, the number of sites segregating is binomially distributed, and obtained 95% confidence intervals on that basis. We tested if the proportion of sites segregating in each compartment is different from the proportion segregating at putatively neutral (here, synonymous) sites using a Fisher exact test, accounting for multiple tests.

We also calculated the fraction of other types of synonymous sites segregating in each sample size of interest (specifically, for T>A variants, and C>Ts not at methylated CpG sites).

### 8. Frequency of mutant alleles in bins of *K* sites

Within each annotation of interest, with an average mutation rate of *u* per site, we construct bins of *k* sites, such that *k*=*U/u*, where *U* is the mean mutation rate of a transition at methylated CpG site in that annotation class. The mean mutation rates are calculated for each mutation type within each annotation, as described in Section 5 above. We then count the fraction of bins in which no such mutations are observed. As an example, for T>A mutations, *k* is on the order of 100 (Supplementary Fig. 11a).

Since each bin can be treated as being comparable to a single neutral methylated CpG site, bins that contain only neutral sites are expected to contain at least one mutation in 99% of bins; this is indeed the case for bins of synonymous sites (Supplementary Fig. 11b).

When considering sites that contain a mixture of neutral and selected sites, bins of *k* sites are no longer as readily comparable to methylated CpG sites, however (Supplementary Fig. 11c). If sites within a bin are under varying degrees of selection, then the mutation count reflects the harmonic mean of the strength of selection acting on individual sites. Specifically, under a deterministic model of mutation-selection balance, if q_i_ is the allele frequency at the *i^th^* site in a bin of *k* sites:

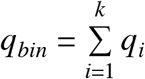

then

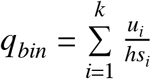

Assuming *u*, = *u* = *U/k*,

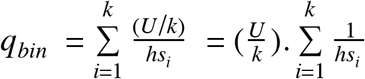

i.e., *q_bin_* is a function of the harmonic mean of *hs* at the *k* sites.

### 9. Forward Simulations

We used a forward simulation framework initially described in Simons et al. (2014), modified in Fuller et al. (2019), and also described in Agarwal, Fuller, and Przeworski (in prep). Briefly, we modeled evolution at a single non-recombining bi-allelic site, which undergoes mutations each generation at rate 2*N_e_u* in a panmictic diploid population of effective population size *N_e_*. Each generation is formed by Wright-Fisher sampling with selection, where fitness is reduced by *hs* in heterozygotes and *s* in homozygotes for the T allele. We fixed the dominance coefficient *h* as 0.5, and we chose one value of the selection coefficient *s* from a log-uniform prior ranging from 10^-7^ to 1 for each simulation (for autosomal mutations with fitness effects in heterozygotes, we only need to specify the compound parameter *hs*; reviewed in ref 31). Given a mutation rate and a demographic model that specifies *N_e_* in each generation, we simulated the evolution of a site forward in time to determine whether the site is segregating in a sample of size *n* at present.

We used *u* = 1.2 × 10^-7^ per site per generation to model CpG>TpG mutation at a methylated CpG site. The simulation framework allows for recurrent mutations, which are expected to arise often at this mutation rate. We also allowed for TpG>CpG back mutations at the rate of 5 × 10^-9^ (calculated from de novo mutation data, as CpG>TpG mutations). To model T>A mutations, we used *u* = 1.2 × 10^-9^ per site per generation, with a back mutation rate of 1.2 × 10^-9^ per site per generation; for C>T mutations not at methylated CpG sites, we used *u* = 0.9 × 10^-8^ per site per generation, with a back mutation rate of 5 × 10^-9^ per site per generation (Supplementary Fig. 1). We note that, since the mutation rate increases with paternal and maternal ages, an implicit assumption is that the distribution of parental ages in the trio data is representative of the parental ages over the evolutionary history of exome samples.

For the demographic model, we relied on the Schiffels-Durbin model for population size changes in Europe over the past ~55,000 generations, preceded by a ~10*N_e_* generation burn-in period of neutral evolution at an initial population size *N_e_* of 14,448 (following ref 34). In the last generation, *i.e*., at present, we sample *n* individuals from the simulated population, to match the size of the sample of interest.

We calculated the probability that a site with the fixed mutation rate *u* is segregating for a given value of *hs* (with *s*=0 under neutrality) as the proportion of simulations with those parameters in which the site is segregating for different sample sizes at present.

In comparing the output of these simulations to data, we considered two scenarios where we may either undercount or overcount segregating CpG sites in the data relative to the simulations. First, because we conditioned on the human reference allele being a CpG in data, we did not count sites where the CpG is the ancestral but not the reference allele. To check how often this is expected to occur, we mimicked this scenario in simulations, sampling a single chromosome at the end of the simulation as the mock haploid reference genome. The proportion of simulations in which CpG is the ancestral but not the reference allele is ~0.1%, i.e., approximately the heterozygosity levels in humans. The second case is that for a subset of the CpG>TpG variants observed at present, the CpG mutation is the reference allele but is not ancestral. To mimic this scenario in our simulations, we simulated a site that starts as TpG (with a mutation rate of 5 × 10^-9^ to CpG, and a back mutation rate ~1.2 × 10^-7^ to TpG) forward in time. Then, as above, we draw a single chromosome from the sample at the end of the simulation and set it as the reference. We obtained the proportion of simulations in which the C allele is the reference, starting from a TpG background. Reassuringly, this occurs in only 0.0014% of simulations. We note that there is in principle a third scenario to consider, in which ApG or GpG sites is ancestral and a C/T polymorphism is found in the sample at present as a result of two mutations, one to T and one to C. Given the various mutation rates involved (all less than 5 × 10^-9^), this double mutation case will be even less likely than the one in which TpG was ancestral. These rare scenarios should not have any substantive effect on our comparison of data to simulations, particularly when we only used such comparisons to examine qualitative trends.

### 10. Inferring selection in simulations

In lieu of calculating the probability that a site segregates for a fixed value of *s*, we can propose *s* from a prior distribution (with *h* fixed at 0.5) and infer the posterior distribution of *hs* for a site with a T allele at 0 copies using a simple Approximate Bayesian Computation (ABC) approach. Specifically, we proposed *s* such that log_10_(s)~U(−7,0); we simulated expected T allele counts under our model for 10 million proposals from the prior. We accepted the subset of the proposed values of *s* where simulations yield 0 copies of the T allele in the sample at present; this set of *s* values is a sample from the posterior distribution of *s* given that the site is monomorphic. We calculated the Bayes odds of s >10^-3^ as the ratio of the posterior odds of s >10^-3^ and the prior odds of s >10^-3^:

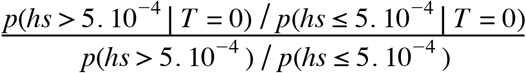

We similarly obtained posterior distributions of *hs* for sites that are segregating at 0, 1-10 copies, or >10 copies, in samples of different sizes, and for three different choices of priors on *s*, namely: s ~ Beta(α=0.001, β=0.1); log(s) ~ N(−6,2); and N_e_s ~ Gamma(k = 0.23, θ = 425/0.23), with *N_e_*=10,000, based on the parameters inferred in Eyre-Walker et al. 2006 (32). These are shown in Supplementary Fig. 3.

### 11. Coalescent Simulations to obtain the length of genealogy of large samples

We simulated the genealogy of a sample of varying sizes using *msprime* (47) under different demographic histories, modifying the standard Schiffels-Durbin model (29) as follows:

a. Demographic history for a sample of Utah residents with Northern and Western European ancestry (CEU) over 55,000 generations, with a recent *N_e_* of 10 million for the past 50 generations, described above.
b. CEU demographic history for 55,000 generations with a recent *N_e_* of 100 million for the past 50 generations.
c. CEU demographic history for 55,000 generations with 5% exponential growth for the past 200 generations.
d. Demographic history for a sample of Yoruba sampled in Nigeria (YRI) ancestry from Schiffels and Durbin 2014, modified with a recent *N_e_* of 10 million for the last 50 generations.
e. A structured sample from two populations that derived from an ancestral population with YRI demographic history 2,000 generations ago, with YRI and CEU demographic histories respectively since, and a recent *N_e_* of 10 million for the last 50 generations in each.

The code for implementing these different demographic models in *msprime* is available as part of the Supplementary Materials. In each case, we recorded the mean genealogy length over 20 iterations.

---

**Supplementary Table 1.**
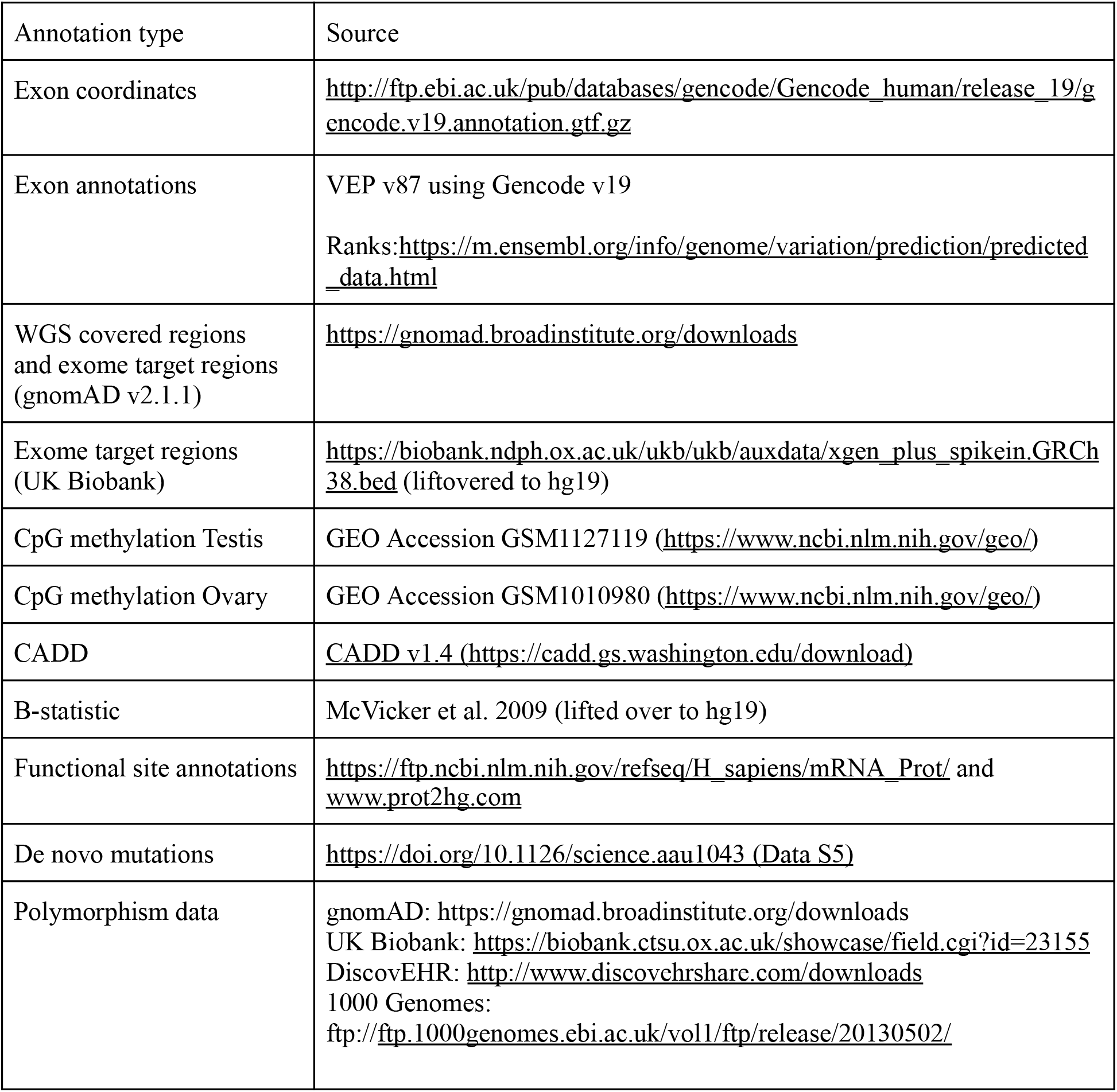

| Annotation type | Source |
| --- | --- |
| Exon coordinates | http://ftp.ebi.ac.uk/pub/databases/gencode/Gencode_human/release_19/gencode.v19.annotation.gtf.gz |
| Exon annotations | VEP v87 using Gencode v19<br>Ranks: https://m.ensembl.org/info/genome/variation/prediction/predicted_data.html |
| WGS covered regions and exome target regions (gnomAD v2.1.1) | https://gnomad.broadinstitute.org/downloads |
| Exome target regions (UK Biobank) | https://biobank.ndph.ox.ac.uk/ukb/auxdata/xgen_plus_spikein.GRCh38.bed (liftovered to hg19) |
| CpG methylation Testis | GEO Accession GSM1127119 (https://www.ncbi.nlm.nih.gov/geo/) |
| CpG methylation Ovary | GEO Accession GSM1010980 (https://www.ncbi.nlm.nih.gov/geo/) |
| CADD | CADD v1.4 (https://cadd.gs.washington.edu/download) |
| B-statistic | McVicker et al. 2009 (lifted over to hg19) |
| Functional site annotations | https://ftp.ncbi.nlm.nih.gov/refseq/H_sapiens/mRNA_Prot/ and www.prot2hg.com |
| De novo mutations | https://doi.org/10.1126/science.aau1043 (Data S5) |
| Polymorphism data | gnomAD: https://gnomad.broadinstitute.org/downloads<br>UK Biobank: https://biobank.ctsu.ox.ac.uk/showcase/field.cgi?id=23155<br>DiscovEHR: http://www.discovehrshare.com/downloads<br>1000 Genomes:<br>ftp://ftp.1000genomes.ebi.ac.uk/vol1/ftp/release/20130502/ |

**Supplementary Fig. 1.**
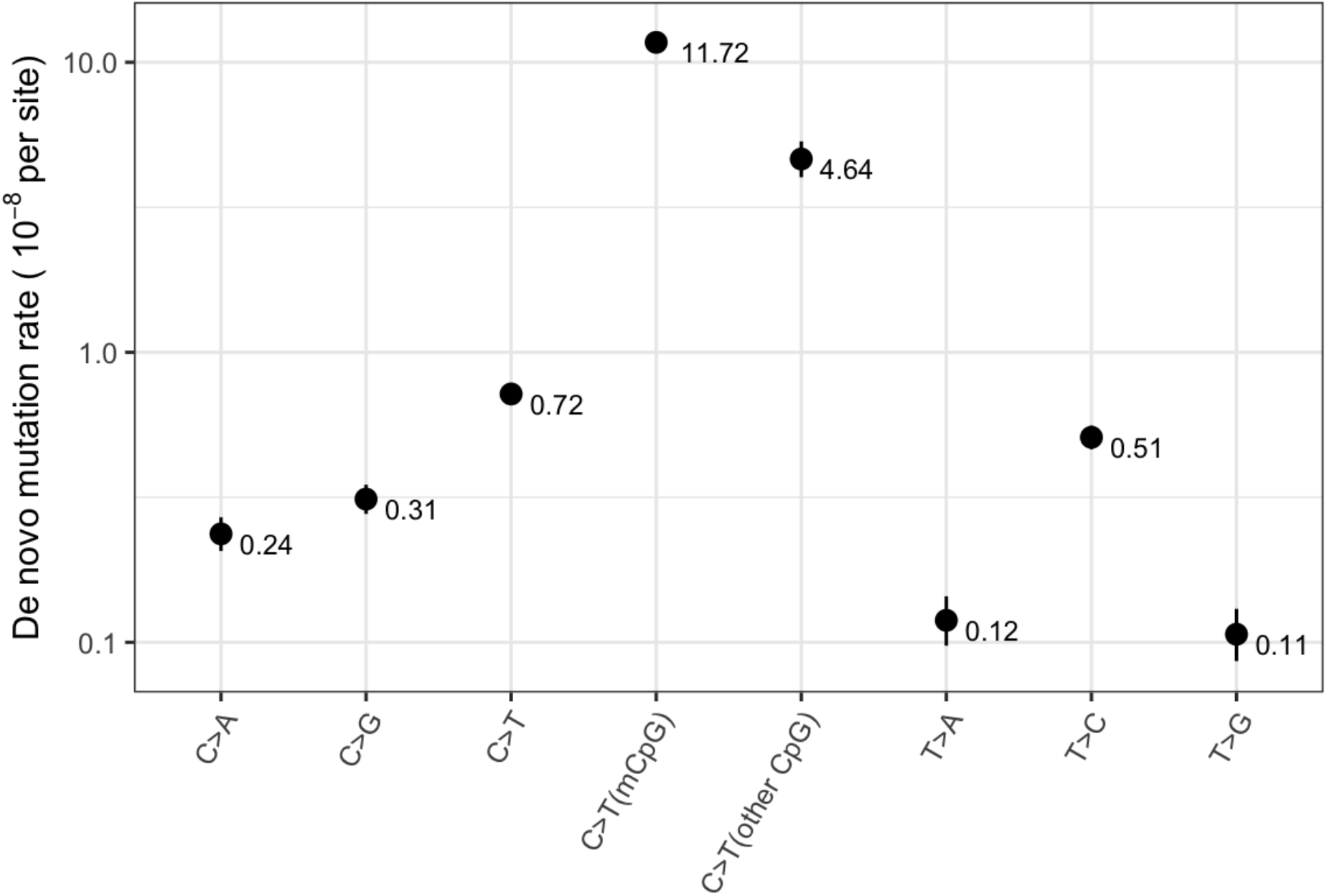
Exonic de novo mutation rates per generation per site estimated from a sample of 2976 parent-offspring trios (data from ref 19), by mutation type. “mCpG” refers to a CpG site with methylation level ≥70% in both testes and ovaries, and “other CpG” to a CpG site with methylation level <70% in either testes or ovaries. Error bars reflect the 95% Poisson confidence interval around mutation counts for each type.

**Supplementary Fig. 2.**
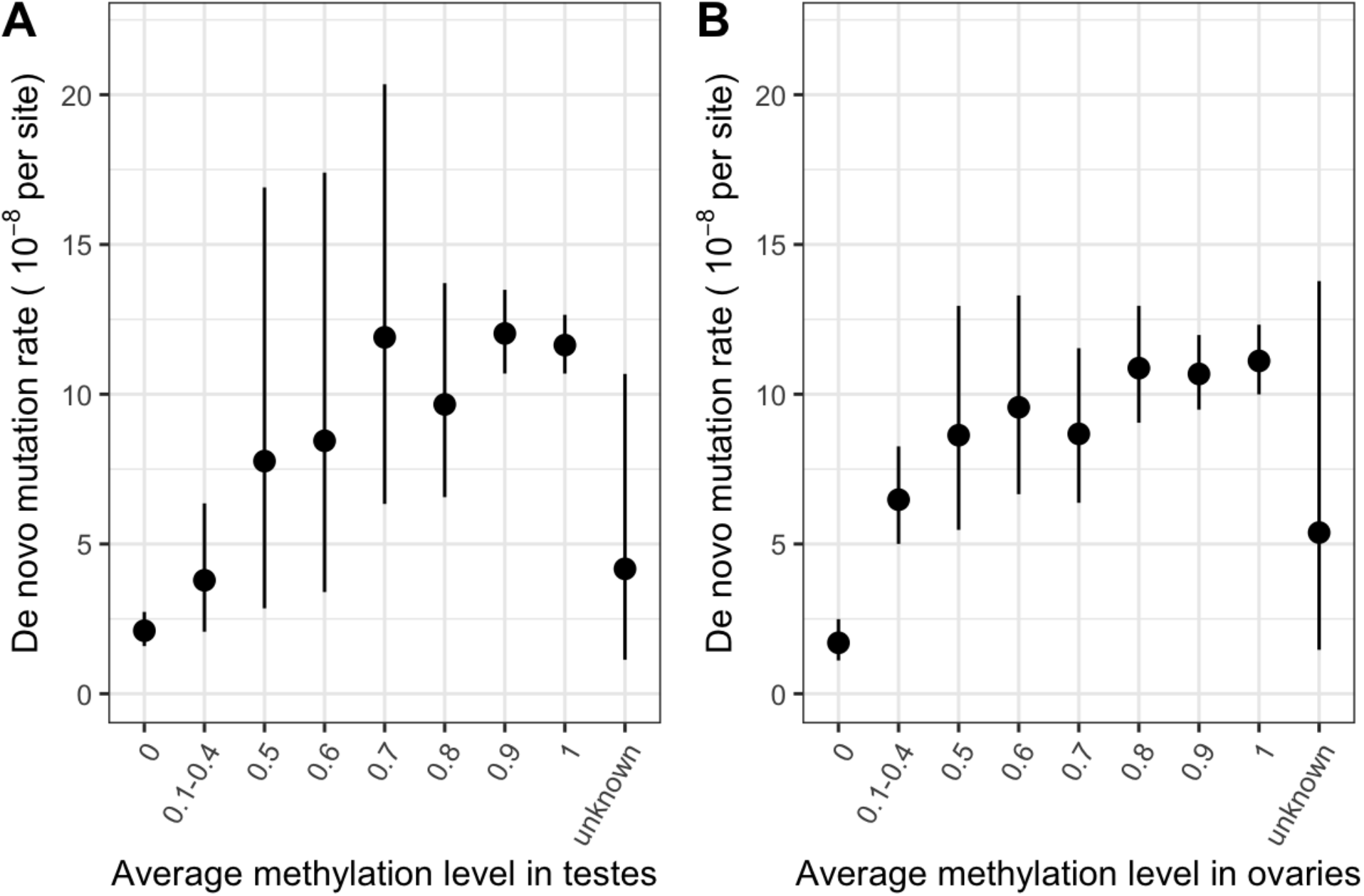
(a) De novo mutation rates in exons in a sample of 2976 parent-offspring trios, by average methylation levels in testes and ovarian tissue. Error bars reflect the 95% Poisson confidence interval around mutation counts in each group (the minimum number of DNMs in each bin is 5).

**Supplementary Fig. 3.**
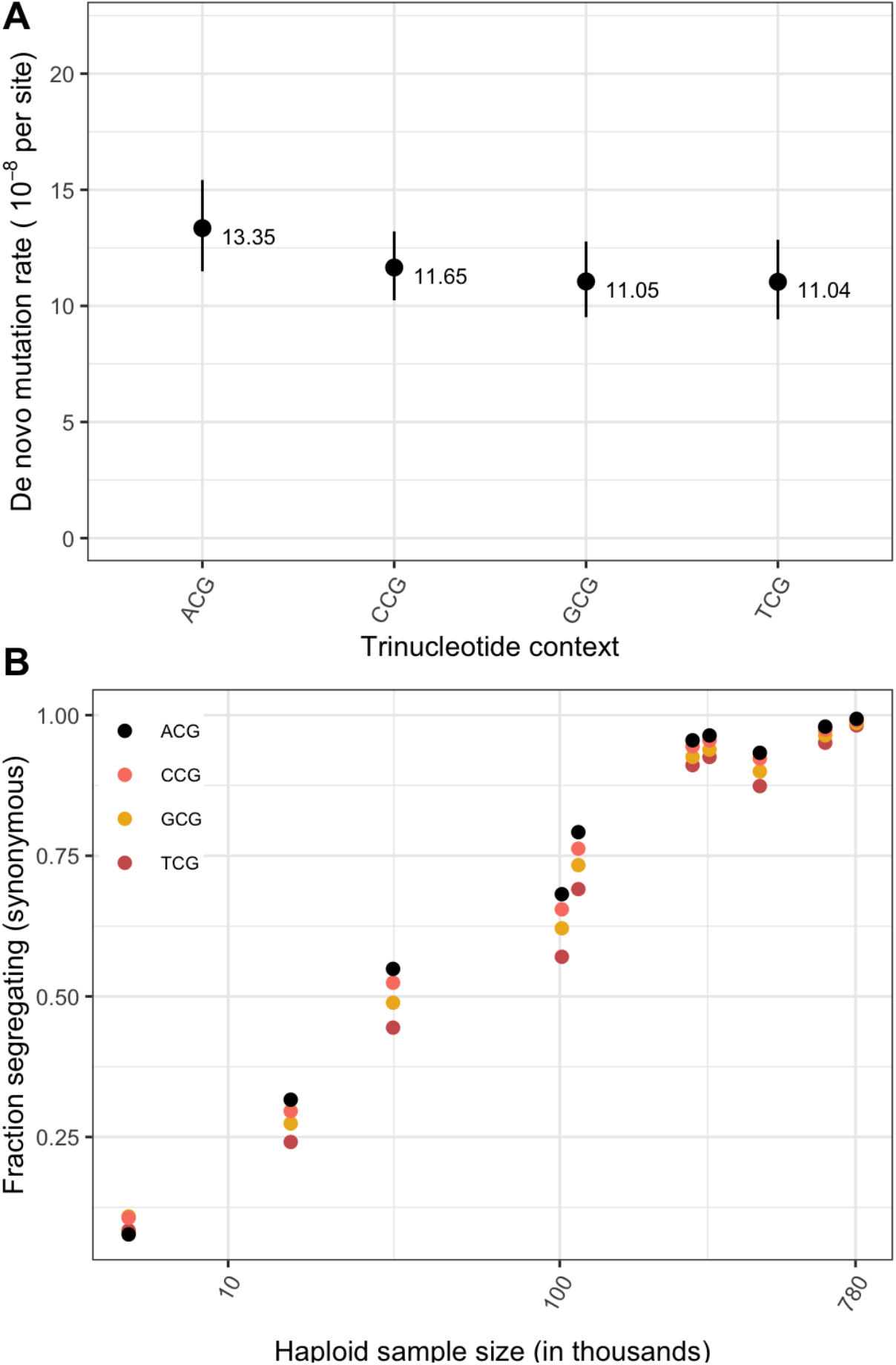
(a) Exonic de novo mutation rates at methylated CpG sites, by trinucleotide context. Error bars reflect the 95% Poisson confidence interval around mutation counts for each context. (b) Fraction of possible synonymous C>T mutations at methylated CpG sites that are observed in a sample of given size, by trinucleotide context.

**Supplementary Fig. 4.**
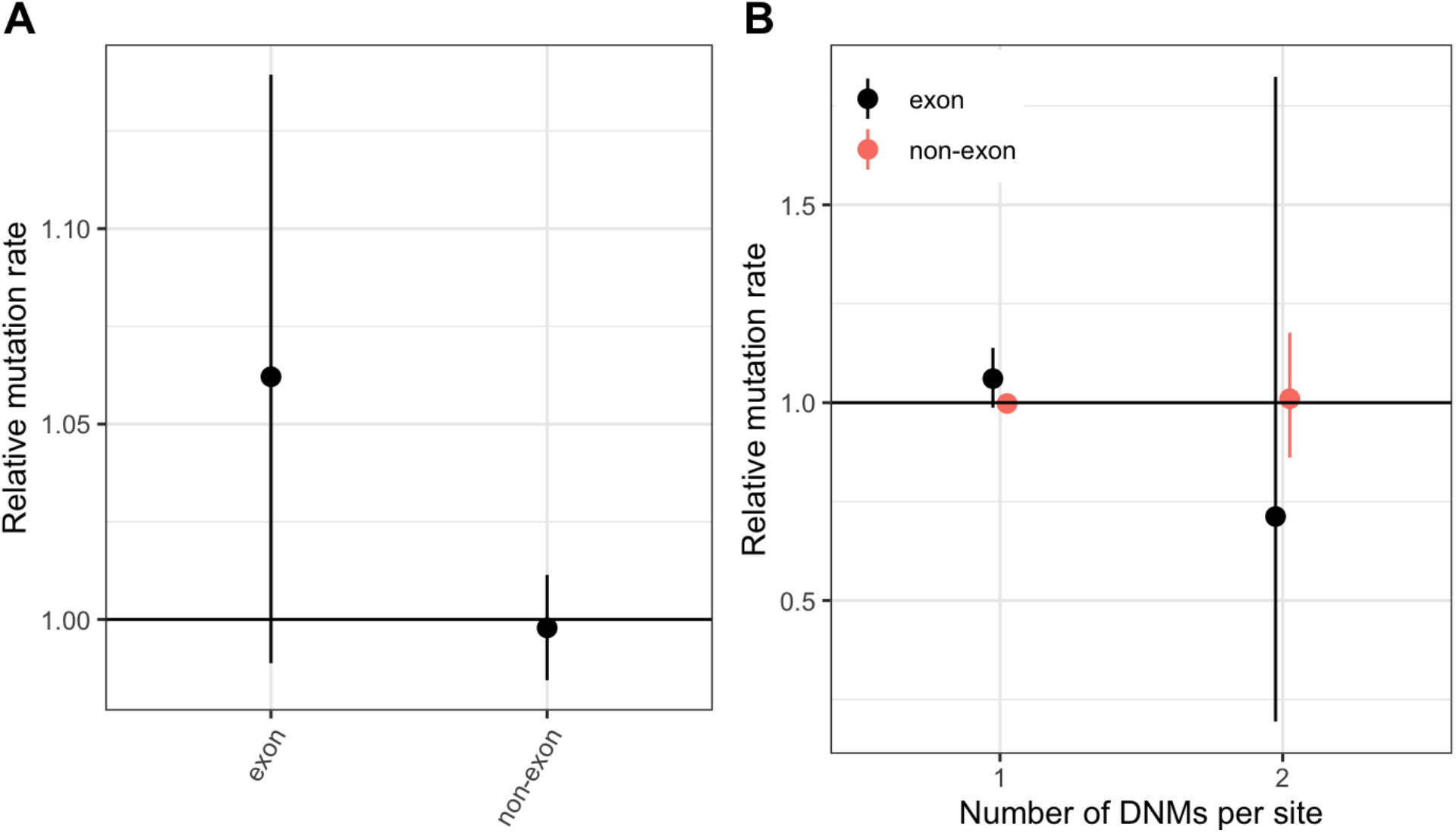
Comparing the distribution of mutation rates in non-exons and exons (a) DNM rates for CpG transitions at methylated sites in exons vs. non-exons, rescaled by the total DNM rate in the genome, with 95% Poisson confidence intervals. (b) The rate of single hits (one DNM at a site) and double hits (two DNMs at a site) in exons vs non-exons, rescaled to the average rate of single and double hits in the genome, with 95% Poisson confidence intervals.

**Supplementary Fig. 5.**
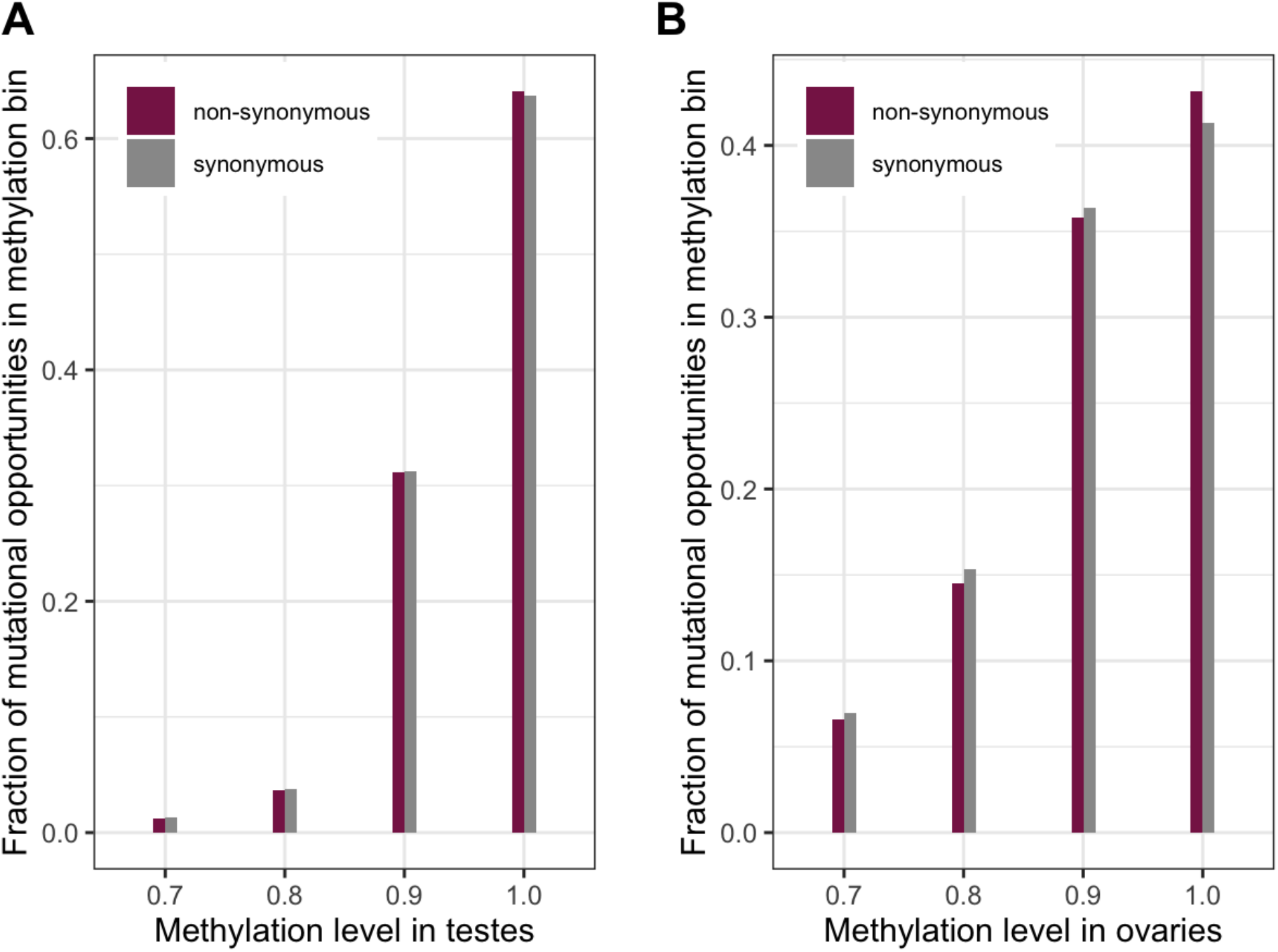
Distribution of methylation levels at synonymous and non-synonymous methylated CpG sites (a) in testes (chi-squared test p-value << 10^-5^) (b) in ovaries (chi-squared test p-value << 10^-5^). The small but significant shift towards higher methylation for non-synonymous sites compared to synonymous ones suggests a small shift towards higher mutation rates at these sites compared to synonymous sites, which should be conservative with regard to identifying non-synonymous sites under selection.

**Supplementary Fig. 6.**
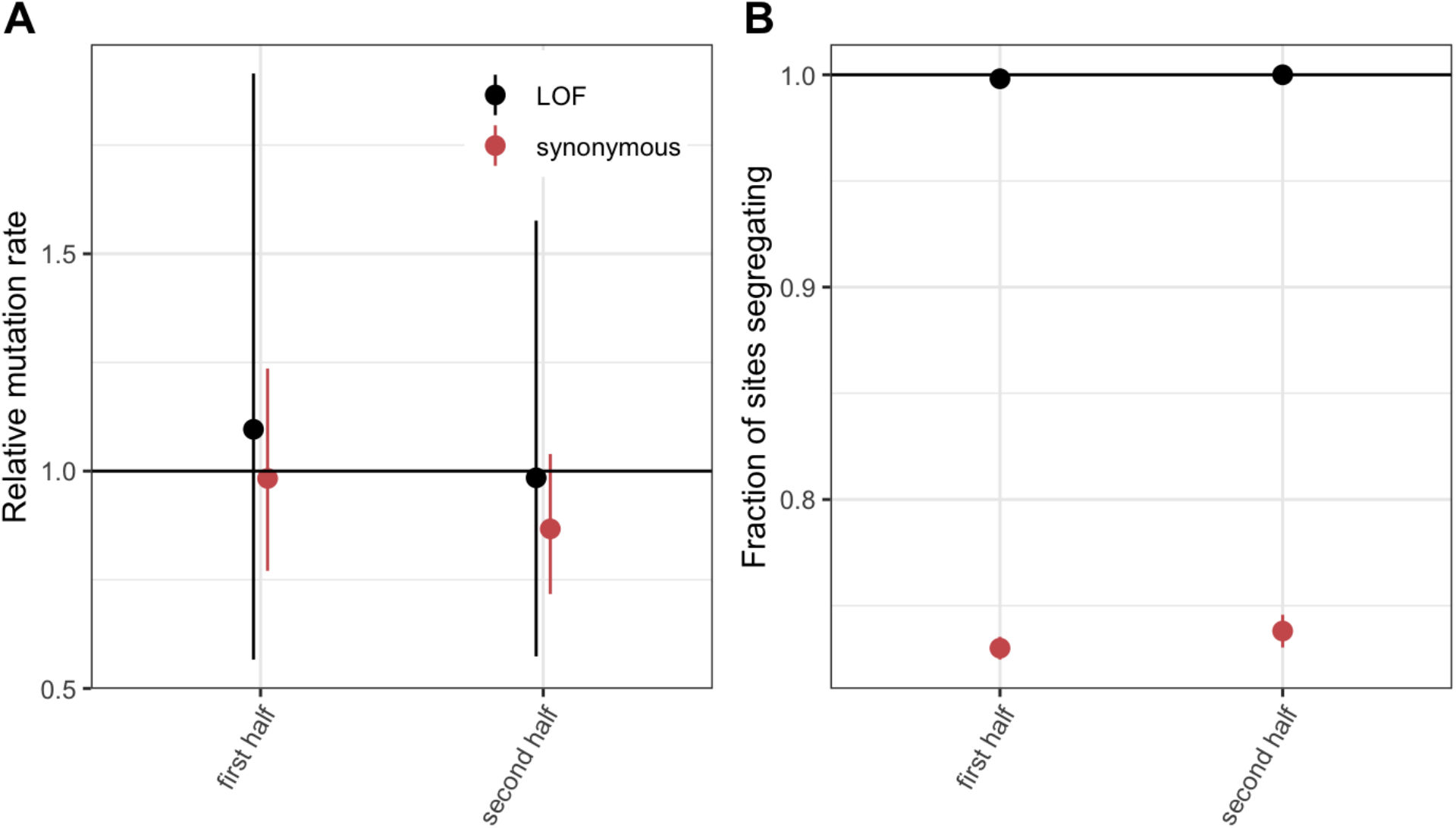
(a) DNM rates for synonymous and LOF CpG transitions at methylated sites in exons that constitute the first vs. second halves of canonical protein coding transcripts, rescaled to the total DNM rate in exons, with 95% Poisson confidence intervals. (b) Fraction of methylated CpG sites that are segregating as a synonymous or LOF C/T polymorphism in exons that constitute the first vs. second halves of canonical protein coding transcripts, relative to the fraction of all synonymous sites segregating. Error bars are 95% confidence intervals assuming the number of segregating sites is binomially distributed (see Methods). LOF variants are defined as stop-gained and splice donor/acceptor variants that do not fall near the end of the transcript, and meet the other criteria to be classified as “high-confidence” loss-of-function in gnomAD (9).

**Supplementary Fig. 7.**
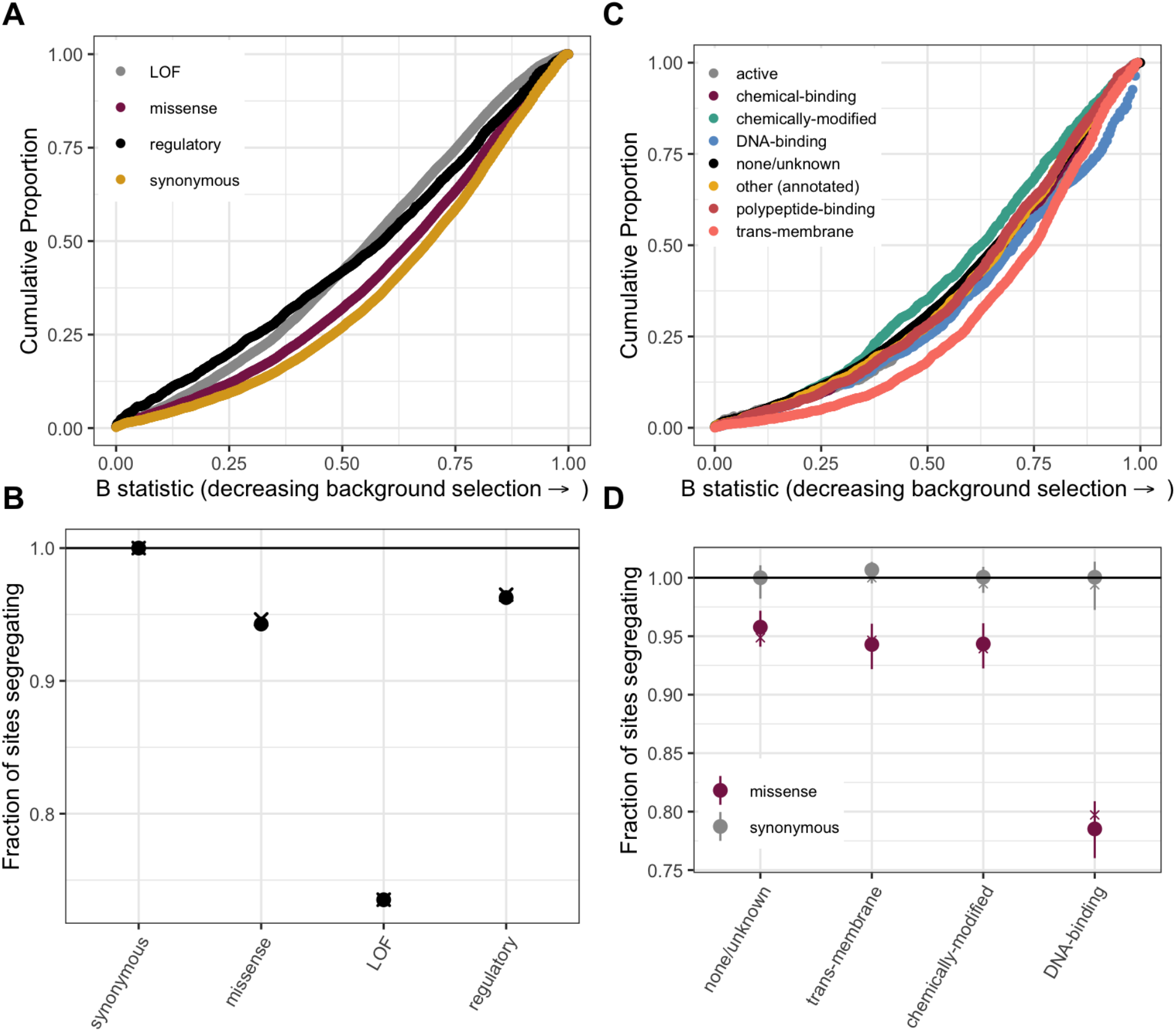
(a) Cumulative distribution of the B-statistic from McVicker et al., 2009 for all possible CpG transitions at methylated sites by annotation class. (b) Fraction of methylated CpG sites that are segregating as a C/T polymorphism in an annotation class, relative to the fraction of synonymous sites segregating, after matching the distribution of the B-statistic across annotations. The fraction segregating without matching for B-statistics (shown in Fig. 3b) is denoted by crosses, to enable comparison. Regulatory variants include non-LOF splice region variants and UTRs. (c) Cumulative distribution of the B-statistic for all possible CpG transitions at methylated sites by functional class. (d) The proportion of synonymous and missense segregating C/T polymorphisms for four functional classes, after matching the distribution of the B-statistic across categories. Error bars are 95% confidence intervals assuming the number of segregating sites is binomially distributed. The fraction segregating without matching for B-statistics (shown in Fig. 3d) is denoted by crosses, to enable comparison.

**Supplementary Fig. 8.**
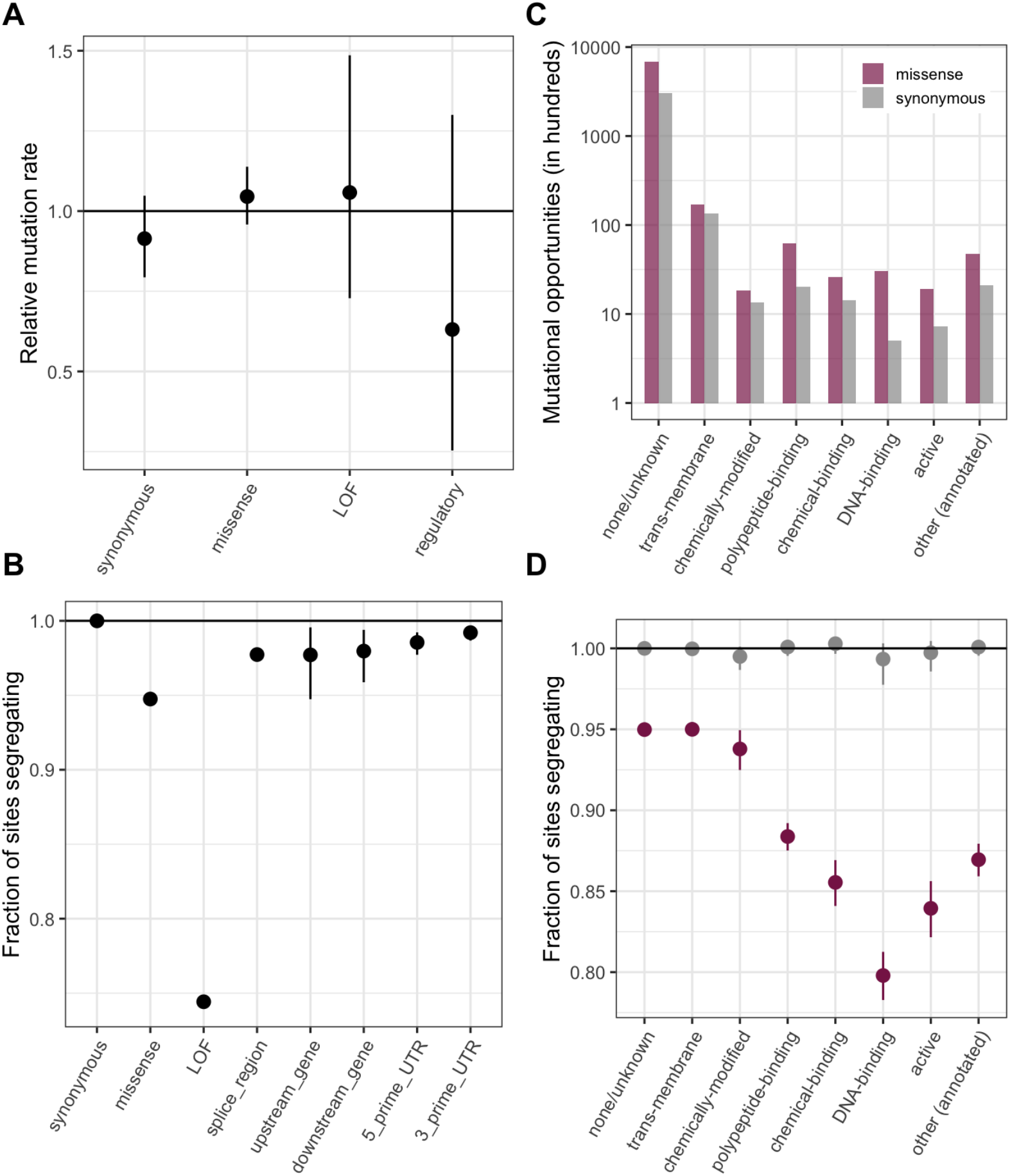
The same analyses as in Fig. 3, but with annotations obtained using the worst consequence in protein coding transcripts by predicted severity, instead of canonical transcripts; the order of preference by which functional sites are assigned to a single category is detailed in Methods. (a) DNM rates for CpG transitions at methylated sites by annotation class, rescaled by the total DNM rate in exons, with 95% Poisson confidence intervals (b) Fraction of methylated CpG sites that are segregating as a C/T polymorphism in an annotation class, relative to the fraction of synonymous sites segregating. Error bars are 95% confidence intervals assuming the number of segregating sites is binomially distributed. LOF variants are defined as stop-gained and splice donor/acceptor variants that do not fall near the end of the transcript, and meet the other criteria to be classified as “high-confidence” loss-of-function in gnomAD. (c) The number of opportunities for synonymous and missense changes involving methylated CpG transitions by the type of functional protein site. (d) The proportion of synonymous and missense segregating C/T polymorphisms in different classes of functional sites.

**Supplementary Fig. 9.**
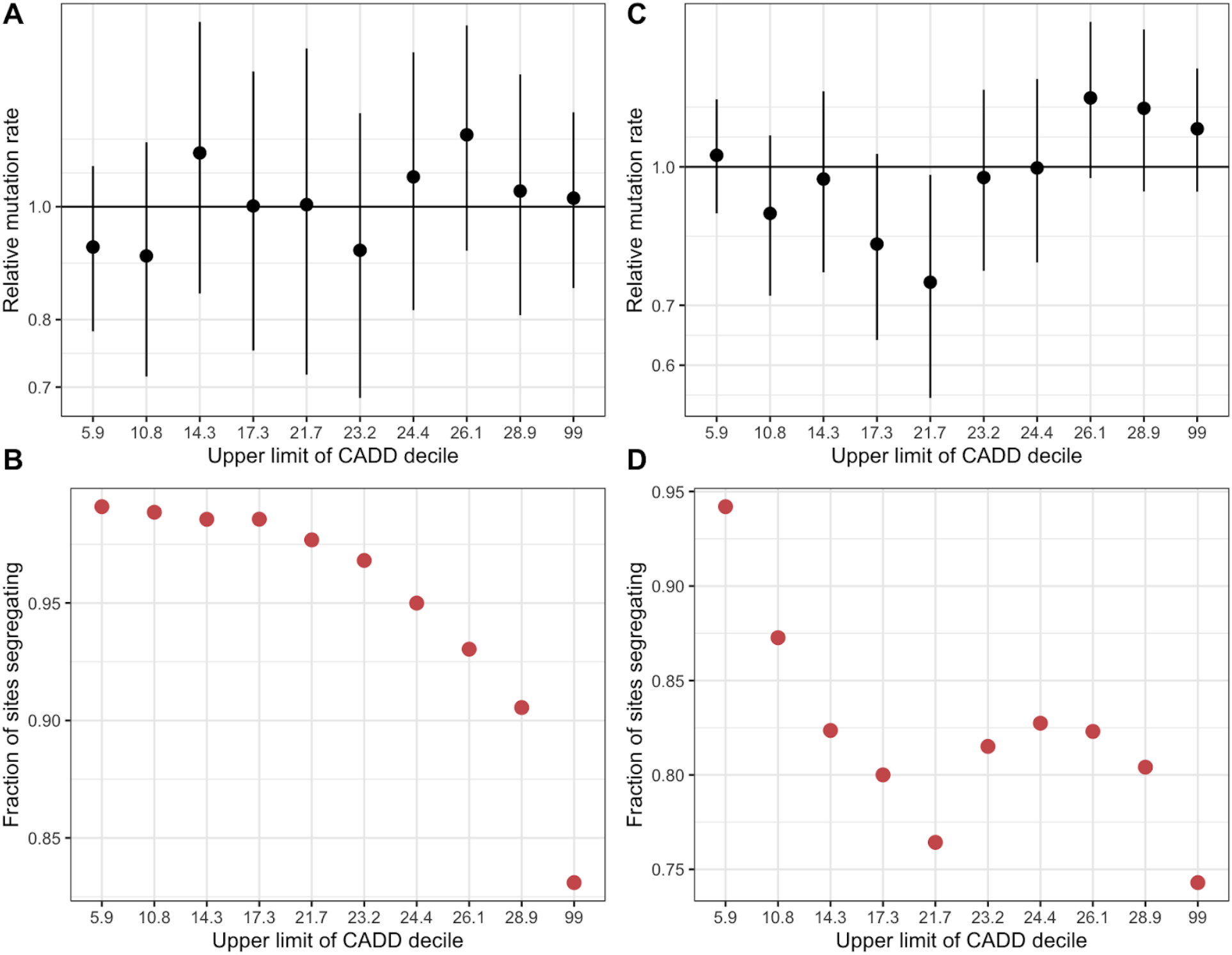
(a) De novo C>T mutation rate at methylated CpGs in deciles of CADD scores in exons, rescaled by the total rate of methylated CpG transitions in exons. Error bars reflect the 95% Poisson confidence interval around mutation counts in each group. (b) Fraction of methylated CpG sites that are segregating as a C/T polymorphism in a CADD score decile, relative to the fraction of synonymous sites segregating. (c) The same as (a) but for C>T mutations at all CpG sites, including unmethylated and less methylated CpGs as well as methylated ones. (d) The same as (b) but for C>T mutations at all CpG sites. Higher CADD scores reflect stronger predicted constraint.

**Supplementary Fig. 10.**
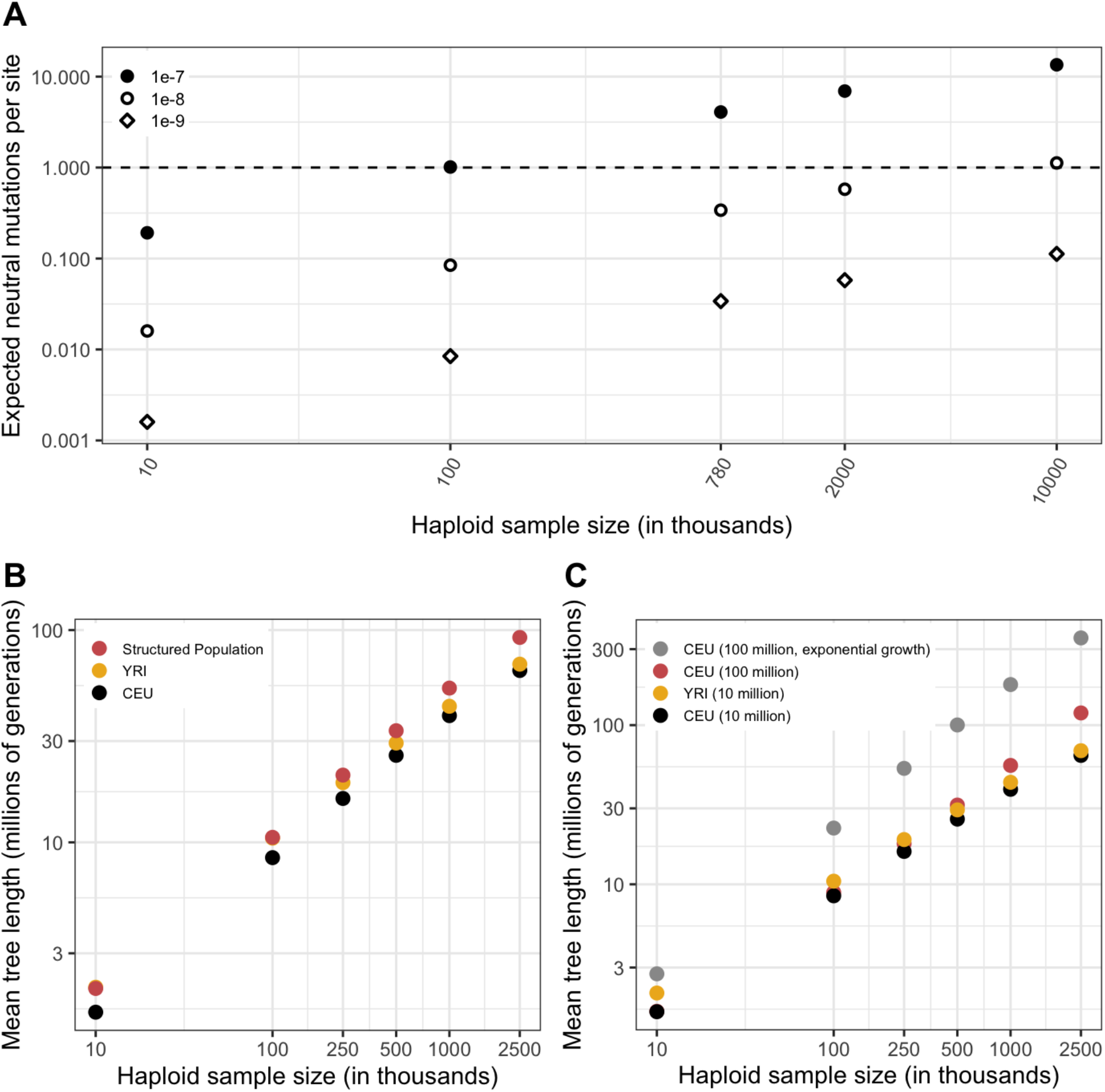
(a) The expected number of neutral mutations in a sample, for three mutation rates, calculated as the expected length of the genealogy (averaged over 20 simulations) for a CEU sample times the mutation rate. (b) A comparison of mean genealogy lengths for four variations on the Schiffels-Durbin demographic models for CEU and YRI populations, namely, YRI demographic history with a recent *N_e_* of 10 million for the last 50 generations, CEU demographic history for 50,000 generations with a recent *N_e_* of 10 million or 100 million, and CEU demographic history with 5% exponential growth for the past 200 generations. (b) A comparison of mean genealogy lengths for samples from YRI and CEU populations, and samples from a structured population derived from an ancestral population 2,000 generations ago.

**Supplementary Fig. 11.**
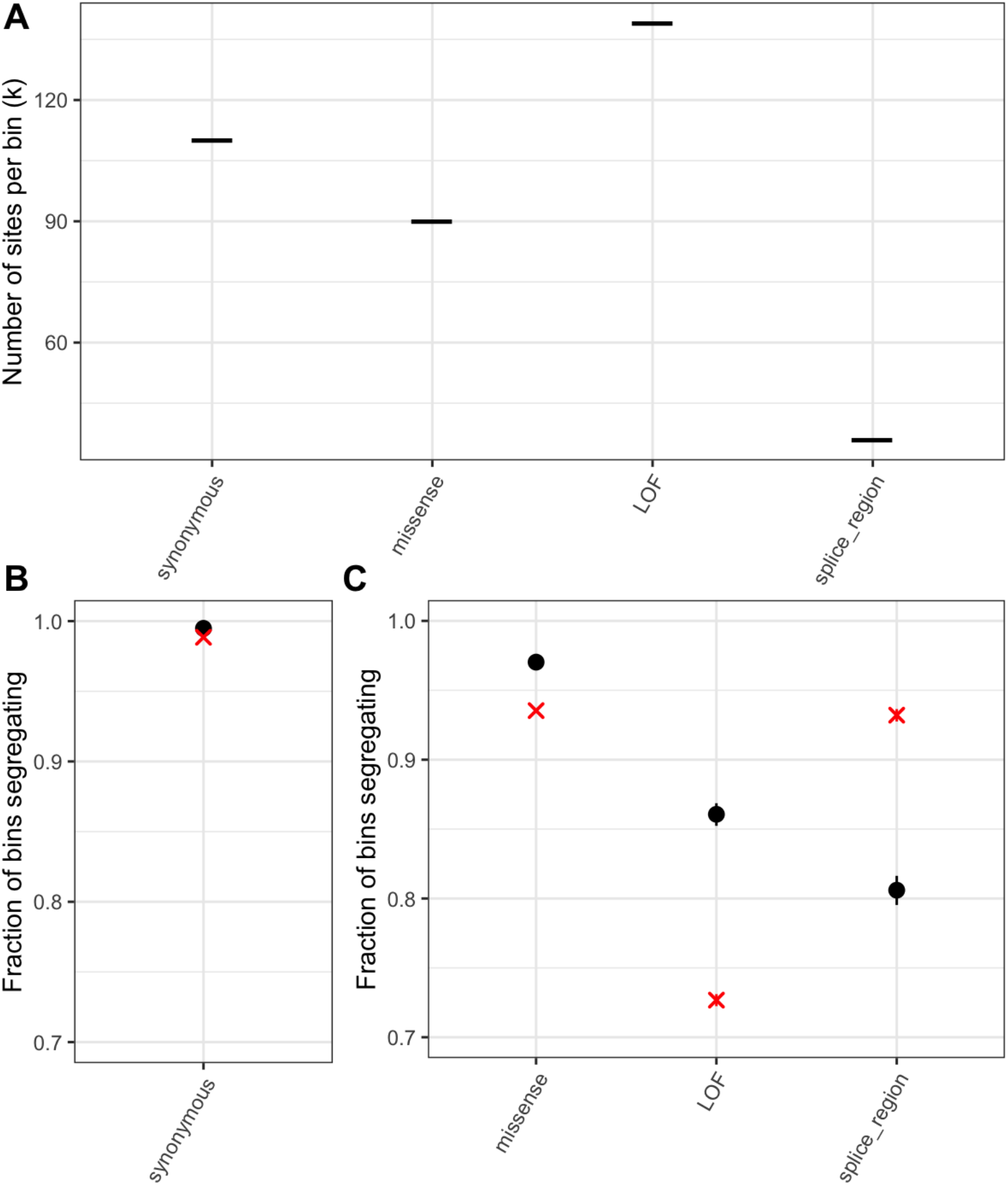
(a) *k*, the number of T sites per bin, such that the average T>A mutation rate per bin is the same as the average transition rate at a single methylated CpG site in that annotation (b) Fraction of bins of synonymous T sites that have at least one T/A polymorphism. A cross is indicated for the corresponding fraction at synonymous methylated CpG sites. As expected if synonymous sites are neutral and the mutation rate for a bin matches that of methylated CpGs, the two fractions are very similar. (c) Fraction of bins that have at least one T/A polymorphism, by non-synonymous annotation. A cross is indicated for the corresponding fraction at methylated CpG sites. Error bars are 95% confidence intervals assuming the number of segregating bins is binomially distributed. For bins including sites under selection the fractions for CpG sites and other mutation types are not expected to match, depending on the extent of variation in mutation rates and fitness effects across sites within a bin (see Methods).

**Supplementary Fig. 12.**
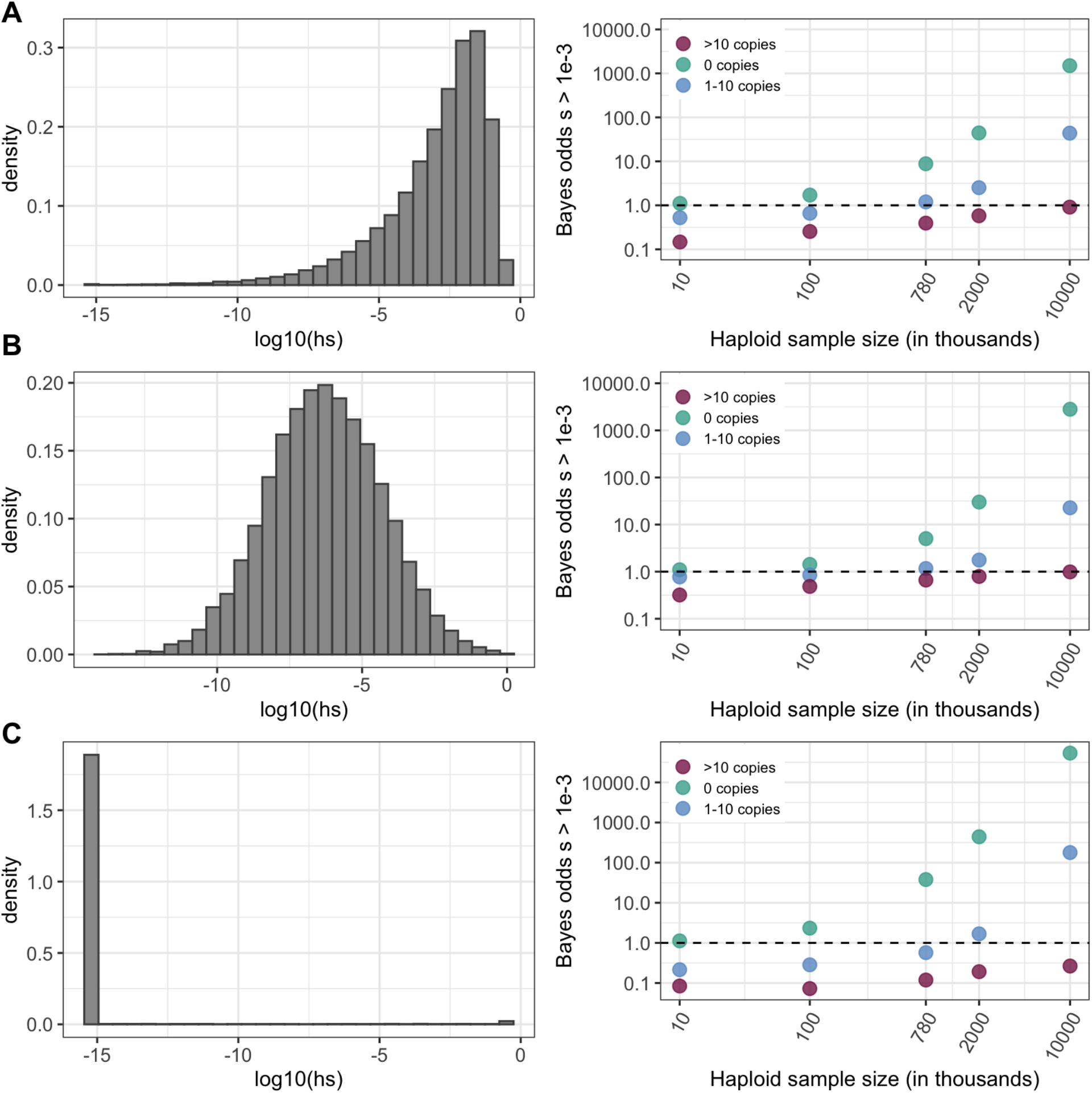
Prior on *hs* (left column) and Bayes odds that *s* > 10^-3^ given that a mutation at a site is observed at 0, 1-10, or >10 copies, for various sample sizes (right column). *h* is fixed at 0.5 (see Methods). The odds are calculated using 10,000 draws from the prior and posterior distributions. (a) *N_e_s* ~ Gamma(shape = 0.23, scale = 425/0.23), with *N_e_*=10,000, the parameters inferred in Eyre-Walker et al. (2006). (b) log(*s*)~N(−6,2) (c) *s*~Beta(alpha=0.001,beta=0.1). Values below 10^-10^ are binned as 10^-10^.

**Supplementary Fig. 13.**
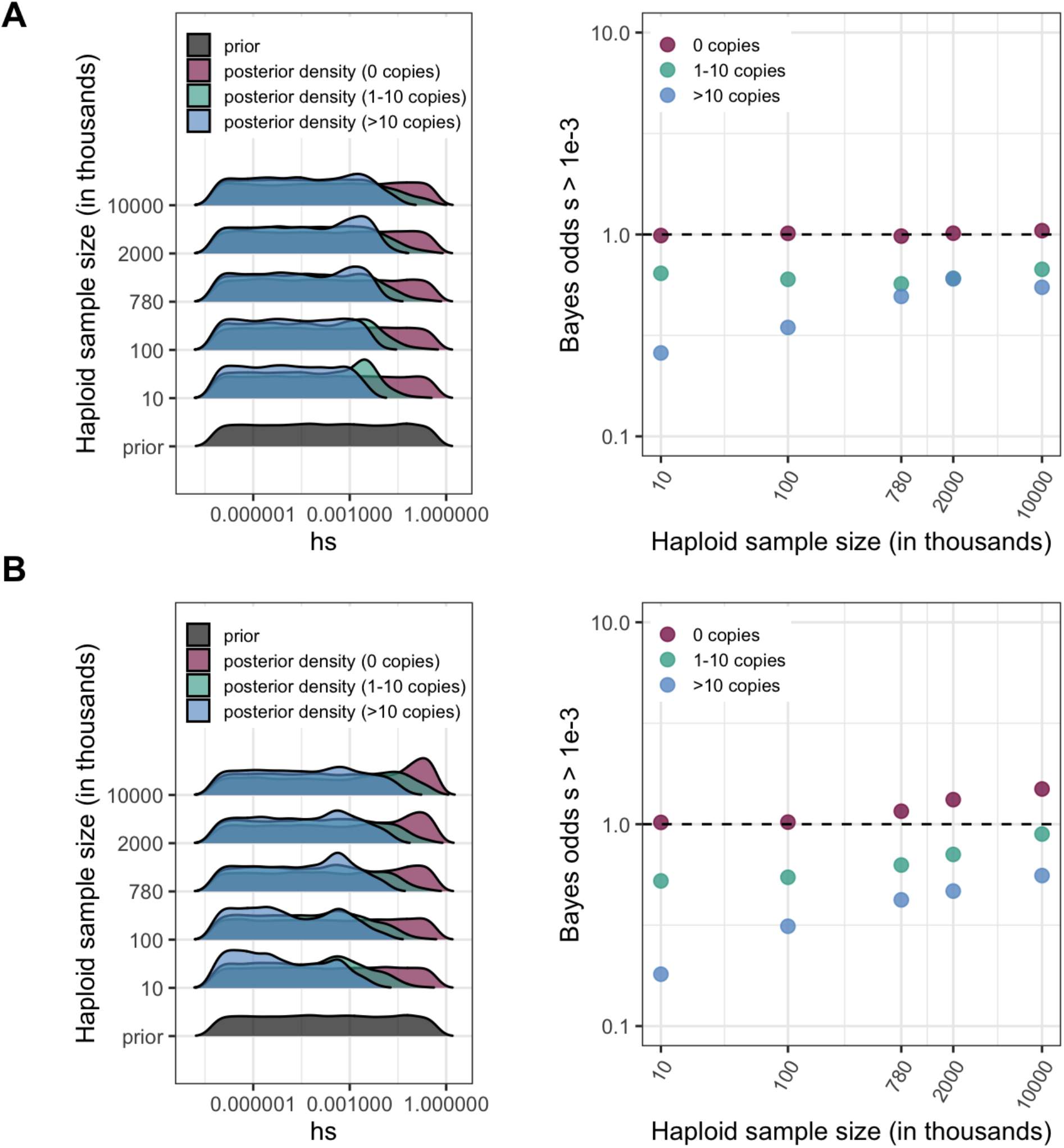
For various sample sizes, prior and posterior log densities for *hs*, and the Bayes odds of *s* > 10^-3^ (and *h*=0.5) for a mutation observed at 0, 1-10, or >10 copies. The prior distribution of *s* is log-uniform over [10^-7^,1]. The odds are calculated from 15,000 draws from the prior and posterior distributions. (a) At a site with mutation rate ~ 10^-9^ (b) At a site with mutation rate ~ 10^-8^.

**Supplementary Fig. 14.**
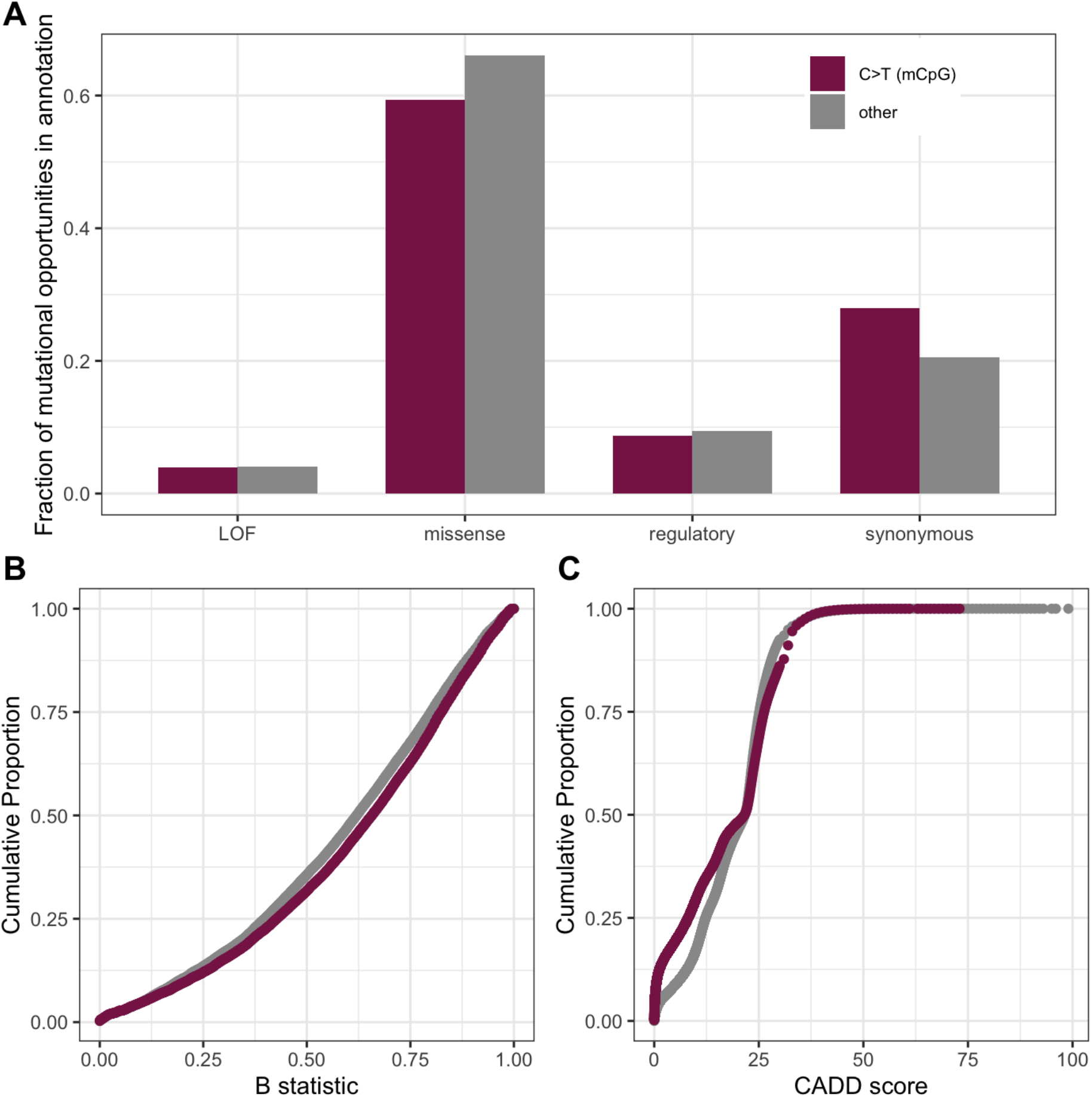
Comparison of measures of deleteriousness at 1.1 million mutational opportunities for methylated CpG (mCpG) transitions vs. 90 million other mutational opportunities in exons (a) Fraction of mutational opportunities for methylated-CpG transitions vs. all other mutational opportunities in exons by their putative functional effect. The difference is statistically significant for missense, regulatory, and synonymous categories (Fisher exact test p-value << 10^-5^) but not for the LOF class (p-value=0.06). (b) Cumulative distribution of the B-statistic from McVicker et al. 2009 for methylated CpG sites vs. all other types of sites in exons (Kolmogorov-Smirnov test p-value << 10^-5^). (c) Distribution of CADD scores at mutational opportunities for methylated-CpG transitions vs. all other mutational opportunities in exons (p-value from a Kolmogorov-Smirnov test << 10^-5^); some skew towards lower values may be expected from the behavior of CADD scores in the presence of mutation rate variation (see Methods). Despite these significant differences, these statistics are overall pretty similar for methylated CpG sites and other mutation types.

## Notes

### Competing Interest Statement

The authors have declared no competing interest.

